# DCLRE1A orchestrates CAG repeat contraction following Cas12a-induced DNA breaks

**DOI:** 10.1101/2025.11.18.689026

**Authors:** Magdalena Dabrowska, Sebastian Siegner, Anna Misiukiewicz, Yauhen Bandaruk, Markus S. Schröder, Matthias Muhar, Susanne Kreutzer, Jacob E. Corn, Marta Olejniczak

## Abstract

Despite their well-defined genetic background, repeat expansion diseases (REDs) still represent an unmet medical need, with no causative therapy offered to patients. The strategy of repeat shortening using genome editing tools is very attractive because a single intervention can result in permanent repair of the disease-causing mutation. However, a limited understanding of DNA repair mechanisms in repetitive sequences complicates the prediction and control of the editing effects. Using a CRISPR interference (CRISPRi) screen, we identified pathways and factors responsible for the repair of staggered cuts generated by Cas12a within CAG repeat tracts. This analysis revealed a central role for interstrand crosslink (ICL) repair factors in mediating CAG repeat contraction, with DCLRE1A emerging as a key effector.

We demonstrated that DCLRE1A recognizes and binds structures generated by Cas12a. Moreover, DCLRE1A interacts with SLX4, promoting the generation of pure contractions, and with POLI, leading to the formation of inverted repeats at the break site during template switching mechanisms. We then used this knowledge to increase the contribution of pure contractions to the pool of editing outcomes using fusions of Cas12a with DNA repair proteins. Our study indicates that Cas12a can be used as an effective tool for generating repeat contractions. Although the mechanisms leading to repeat shortening are complex, understanding them can help researchers develop more precise therapeutic strategies with greater control of the editing process.

## INTRODUCTION

Microsatellite expansions are responsible for almost 50 human diseases [1]. Among these diseases are polyglutamine (polyQ) diseases, which are caused by the expansion of CAG trinucleotide repeats in coding regions of unrelated genes and include Huntington’s disease (HD), dentatorubral–pallidoluysian atrophy (DRPLA), spinal–bulbar muscular atrophy (SBMA), and several spinocerebellar ataxias (SCAs) [2]. PolyQ diseases are incurable and are characterized by progressive neurodegeneration and severe motor and cognitive impairments. As they are monogenic in nature, these diseases are ideal targets for gene therapies. For each polyQ disease, the length of the CAG sequence characteristic of the mutant variants has been determined, and it usually reaches > 40 triplet units. However, due to somatic instability, the length of CAG repeats may increase in some tissues during a patient’s lifetime. Recent studies have indicated that somatic expansions may contribute to the selective vulnerability and degeneration of certain types of neurons [3][4]. Disturbances in DNA repair, especially in the mismatch repair pathway, are considered the leading cause of repeat instability [5]. A genome-wide association study (GWAS) identified MSH3, MLH1, FAN1, LIG1, PMS1, and PMS2 as genetic modifiers of HD onset [6]. This knowledge was used to propose a new therapeutic strategy to limit somatic expansion and halt the progression of HD by modifying the levels of DNA repair proteins. An alternative strategy is to shorten CAG repeats to a nonpathological length using genome editing tools, such as zinc-finger nucleases, transcription activator-like effector nucleases (TALENs), and the CRISPR-Cas9 system. Studies in yeast models, human cells, and mice have shown that the induction of double-strand breaks (DSBs) or several single-stranded DNA nicks within the repeat tract leads to the shortening of the CAG repeats [7][8][9][10][11][12]. However, the DNA repair mechanisms involved in this process are poorly understood. This process is of particular interest because the expanded CAG/CTG sequence forms stable secondary structures that can affect the binding of DNA repair factors, the initiation and kinetics of DNA strand resection, and other steps of DNA repair [13][14][10]. The choice of repair pathway depends on many factors, including the structure of the DNA ends, the degree of microhomology (MH), the availability of repair proteins, and the cell cycle status [15]. Knowledge of the mechanisms involved in endonuclease-induced microsatellite instability may be used to predict and control editing results, e.g., by promoting MH-mediated repeat contractions.

Shortening of the CAG repeats using the CRISPR-Cas9 system and its nickase version occurs through the recognition of a noncanonical NAG protospacer adjacent motif (PAM) (i.e., CAG) and the generation of DNA breaks within the repeats. This method has the risk of off-target gene editing since gRNAs composed of pure CAG sequences may recognize many complementary sequences in the genome. A way to increase specificity may be to use the type V CRISPR-Cas12a nuclease, which recognizes T-rich PAMs and introduces staggered cuts located distal to the PAM sequence [16]. The CRISPR-Cas9 system and Cas12a-based platforms have more distinct features. A one-part guide RNA, called CRISPR RNA (crRNA), is much shorter than the Cas9 gRNA, and Cas12a proteins are typically smaller than the SpCas9 nuclease, simplifying their vectorization [17]. Cas12a showed little or no tolerance for mismatches in mammalian cells and high target specificity in vivo [18]. Therefore, it is considered a more specific alternative to SpCas9 [19]. Cas12a leaves ∼5 nt 5’ overhangs that attract different DNA repair proteins than the blunt ends generated by Cas9. This process may affect the editing results, as Cas12a has been shown to induce mainly deletions and deletions combined with small insertions and not DNA insertions, which are routinely observed for Cas9 [20].

Two polyQ genes, *ATN1* and *ATXN3*, contain an *Acidaminococcus* sp. Cas12a (AsCas12a)-specific TTTN PAM in sequences flanking CAG repeats. Fortunately, new Cas12a variants have been engineered to recognize a broader range of PAMs (TTN, TNN, and VTTV), allowing targeting across all known polyQ disease loci [19], [21]. This approach allows the crRNA to be anchored to specific sequences of these genes and generate DNA breaks directly within the CAG repeats. As CRISPR-Cas12a has not yet been used to edit expanded CAG repeats in human cells, it is necessary to explore the possibilities and limitations of this repeat shortening method.

Here, we demonstrate that the repair of AsCas12a-induced DNA breaks leads to the gradual shortening of CAG repeats. However, alongside pure MH-based CAG repeat contractions, we observed characteristic insertions that changed the reading frame of the protein-coding mRNA. Using a dedicated CRISPR interference (CRISPRi) screen and biochemical approaches, we identified DNA repair pathways and candidate genes involved in the induction of specific editing outcomes. We have shown that DCLRE1A recognizes and physically binds the structures generated by Cas12a in cells. It cooperates with SLX4, increasing the number of pure MH-based contracted variants. Moreover, we captured a new interaction between DCLRE1A and POLI. Their concert actions during template switching mechanisms led to the formation of inverted repeats at the break site, inhibiting the gradual shortening of the repeats. Finally, we created hybrids composed of Cas12a and DNA repair proteins to control the process of CAG repeat contraction and increase pure repeat shortening. Our proof-of-concept study showed that Cas12a can be used for the controlled shortening of mutated CAG repeat tracts, which holds significant therapeutic potential for currently incurable disorders.

## RESULTS

### AsCas12a and SpCas9 generate different patterns of CAG repeat editing outcomes

We generated a homozygous HEK293T cell line containing 35 CAG repeats in the *ATN1* locus (HEK293 T_35CAG_ATN1) to explore CAG repeat instability following AsCas12a-induced DSBs. Although 35 CAG repeats fall in the intermediate range between normal and pathological lengths in many polyQ diseases, these repeats are considered unstable and tend to expand. For Cas12a, we used an RNP complex, composed of AsCas12a and g14, with a 5’-TTTC-3’ PAM located downstream of the CAG tract to introduce a staggered cut within the CAG repeats. To compare Cas9 with Cas12a, we utilized two Cas9 gRNAs (g10 and g11) that recognize 5’-TGG-3’ PAMs and generate cleavages within glutamine-encoding CAG/CAA triplets located 5’ upstream of pure CAG repeats (Figure 1A). Cells were transfected with RNP complexes, and 24 h later, the region of interest was amplified and subjected to sequencing. CAG repeat contraction upon Cas12a- and Cas9-mediated cleavage was confirmed by agarose gel electrophoresis (Figure 1B) and by the NGS results (Figure 1C-1D, Supplementary Figure 1A). Contraction events resulting from polymerase slippage during the PCR amplification of a long tract of CAG repeats were also observed in the unedited sample (Supplementary Figure 1B). The sequencing results were divided into six categories: ‘35 CAG’ corresponding to preserving the CAG-35 output sequence, ‘Del CAG’ and ‘Del CAG&CAA,’ indicating the deletion of CAG repeats or the deletion of CAG repeats and CAA interruption, respectively (Figure 1E). Other events included ‘Indel CAG& flanking region,’ indicating that the deletion of the CAG repeat extended to sequences flanking the repetitive tract, sometimes accompanied by small insertion events. Moreover, we distinguished the ‘Indel CAG’ and ‘Ins CAG’ categories corresponding to several nucleotide insertions/deletions within the CAG repeat tract and expansions of the CAG repeats, respectively (Figure 1E).

**Figure 1.**
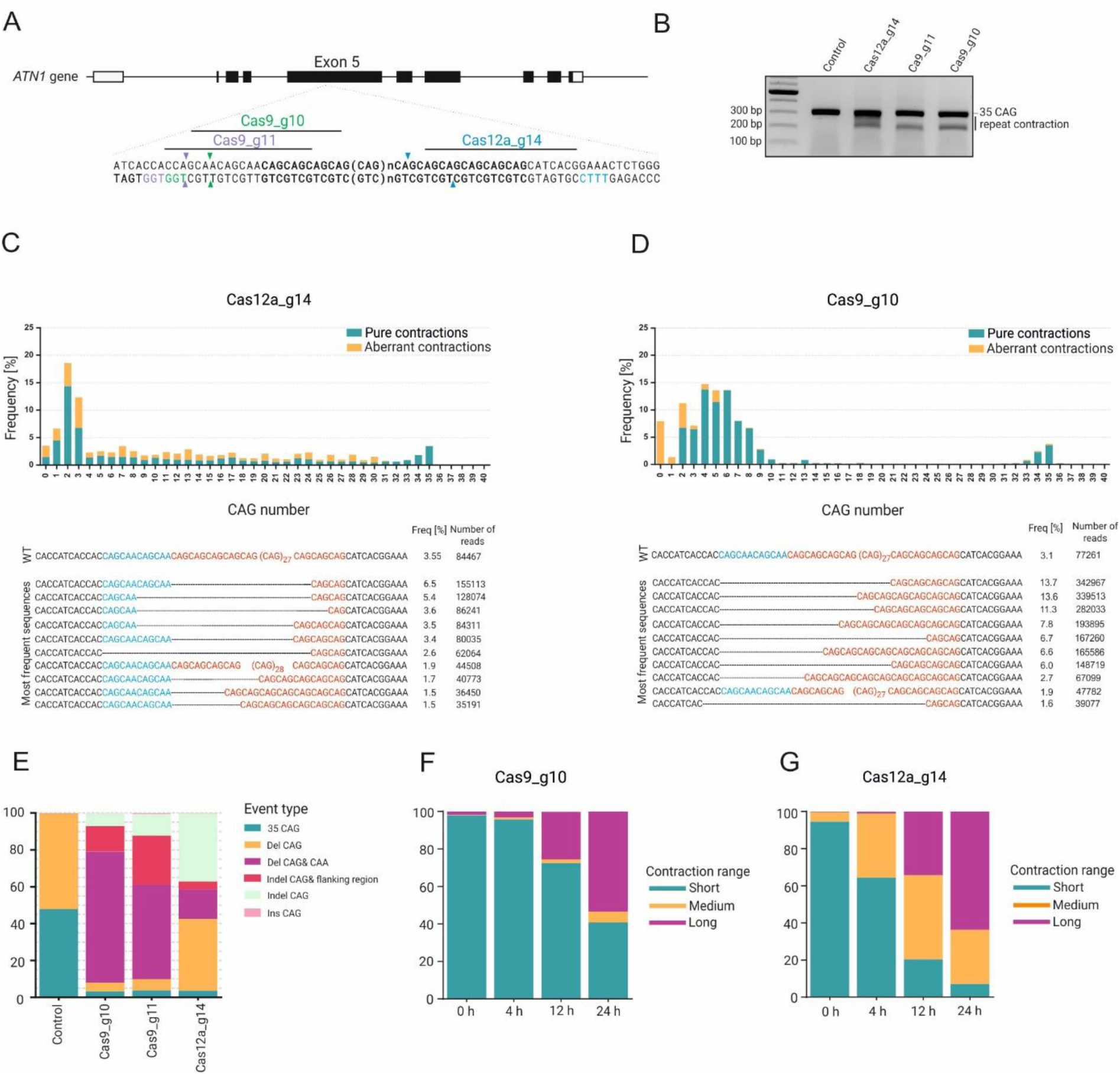
Genome editing strategy for shortening CAG repeats in the ATN1 gene. **A.** The polymorphic CAG repeat tract with adjacent CAA interruptions is located in exon 5 of the ATN1 gene. Cas9 guide RNAs (g10 and g11) were designed to recognize glutamine-encoding CAG/CAA triplets, whereas Cas12a_g14 was intended to target the CAG repeat tract directly. The appropriate PAM sequences are highlighted in green, violet, and blue for the RNP complexes composed of Cas9_g10, Cas9_g11, and Cas12a_g14, respectively. **B.** Agarose gel showing the contraction of CAG repeats after Cas9- and Cas12a-mediated cleavage. **C–D**. Graphs showing the frequency of NGS reads with the indicated number of CAG repeats. The top ten sequences are presented below the graph. The CAG/CAA triplets located 5’ upstream of pure CAG repeats and the pure CAG repeat tract are highlighted in blue and red, respectively. The deletions of the CAG repeats after Cas9- and Cas12a-mediated cleavage are represented as dashed lines. **E.** The genome editing outcomes of edited HEK293T_35CAG_ATN1 cells were categorized into six main groups. **F–G.** The contraction range following Cas9_g10- and Cas12a_g14-mediated cleavage, as determined by capillary gel electrophoresis. Cells were collected at 0, 4, 12, and 24 h after the start of the editing process.

For both CRISPR‒Cas systems, we observed a similar editing efficiency, as unedited variants with 35 CAG repeats represented only ∼4% of all the sequences. However, CAG contraction patterns varied significantly depending on the nuclease used. The number of CAG repeats gradually decreased in cells treated with AsCas12a, with a dominant fraction of variants having 0–3 CAG repeats (Figure 1C). Cas9 generated mainly sequence variants with 2–10 CAG repeats, accounting for 84% of the reads (Figure 1D, Supplementary Figure 1A). ‘Del CAG&CAA’ was the most frequent variant, constituting ∼71% and ∼51% of the reads for Cas9_g10 and Cas9_g11, respectively (Figure 1E). Thus, the number of CAG repeats was shortened mainly within the polyQ-encoding sequence, extending the deletion into the left flank, which is composed of a CAA interruption that also encodes glutamine in the ATN1 transcript. ‘Del CAG’ consists of approximately 4.8% and 6% for Cas9_g10 and Cas9_g11, respectively. ‘Del CAG&CAA’ and ‘Del CAG’ will later be called “pure contractions” in this work (Supplementary Figure 1C). ‘Indel CAG& flanking region’ was present in 13.8% and 26.7% of the reads for Cas9_g10 and Cas9_g11, respectively (Figure 1E). Another group was ‘Indel CAG’, which accounted for approximately 10% of the events in both cases. Unlike those of Cas9, a significant fraction of AsCas12a products were shortened by disrupting the polyQ-encoding sequence (42%) (‘Indel CAG& flanking region’ and ‘Indel CAG’). These will be referred to as aberrant contractions below. After Cas12a-mediated cleavage, three main fractions of events were observed: ‘Del CAG’, which accounted for 39% of the reads, ‘Indel CAG’ (36.9%), and ‘DelCAG&CAA’ (16.1%) (Figure 1E). A characteristic event in the ‘Indel CAG’ group was inverted repeats, a CTG mutant variant (Supplementary Figure 1D).

Next, we analyzed the dynamics of the repeat shortening process using hTERT RPE-1 cells previously infected with a lentivirus carrying 40 CAG repeats (RPE-1_40CAG). The cells were transfected with AsCas12a or SpCas9 RNPs, and the repair products were characterized at various time points (from 0 h to 24 h) by capillary electrophoresis and amplicon sequencing. Repair of DNA breaks induced by Cas9_g10 resulted in long deletions (approx. 30 CAG repeats), which appeared 4 h after RNP transfection and increased over time (Figure 1F, Supplementary Figure 2A). In contrast, Cas12a_g14 shortened the CAG sequence by approximately 5–15 units and progressed until 24 h (Figure 1G, Supplementary Figure 2B). Pure contracted products accounted for ∼80% of all the reads at 4 h, and their number decreased to 54.6% and 22.5% at 12 and 24 h, respectively (Supplementary Figure 3A). Simultaneously, products containing inverted repeats at the cleavage site appeared over time (Supplementary Figure 3B). This process resulted in the loss of complementarity of Cas12a_g14 to the target sequence, thus stopping the editing process.

These data indicate that the mechanism of repair of Cas12a-induced DNA breaks is complex and dynamic and leads to the generation of both (i) therapeutically desirable CAG repeat contractions and (ii) indel mutations that result in gene knockout by a frameshift. The number of these undesirable variants increases with time.

### A CRISPR interference screen reveals factors involved in the repair of Cas12a-induced DNA breaks within CAG repeats

To better understand the complex mechanism after Cas12a-induced breaks, we performed a pooled CRISPR interference (CRISPRi) screen. CRISPRi utilizes a catalytically inactive SpCas9 (dCas9-KRAB) fusion protein and a gRNA library to inhibit the transcription of target genes and analyze specific readouts in a high-throughput manner [22]. As a model, we used hTERT RPE-1 cells with stable expression of dCas9-KRAB. The cells were transduced with lentiviral screening vectors that link whole genome library (WGL) gRNA expression to a nearby ‘‘target region’’ composed of 40 CAG repeats and short flanking sequences of the *ATN1* gene (Figure 2A). DSBs within CAG repeats were generated using the Cas12a_g14 RNP complex, and DNA was isolated 2 days later. General screen performance was evaluated by confirming the dropout of many essential genes (Supplementary Figure 4A). Therefore, as a next step, we analyzed the gRNA sequence and the length of the CAG repeat-containing sequence (target sequence) to identify specific genes from WGL whose knockdown might change the contraction of CAG repeats. For the analysis of the CAG repeat length, the R2 read containing the target sequence was analyzed for the presence of an anchor sequence up- and downstream of the CAG repeat region (Supplementary Figure 4B). The sequence length between the anchor sequences was calculated, and the sequences were then sorted into bins (Supplementary Figure 4C). We used nontargeting gRNAs to create a reference distribution (Supplementary Figure 4C) and looked for absolute differences in genes compared with the nontargeting distribution. After confirming a good correlation of this metric between replicates (Supplementary Figure 4D), we combined replicates and looked for candidate genes for which three sgRNAs showed a similar contraction pattern that significantly differed from the nontargeting distribution using the G test [23]. We identified genes involved in diverse cellular processes, including those related to DNA repair (Figure 2B, Supplementary Table 1, Supplementary Figure 5). Among them were FANCD2, which forms a complex with FANCI and recognizes branched DNA structures, and UBE2T, which is responsible for its ubiquitination (Figure 2B) [24]. These genes are related to interstrand crosslink (ICL) repair, Fanconi anemia (FA), and homologous recombination (HR) [25]. In addition, other genes that also changed CAG contraction pattern, such as DCLRE1A, PMS2, and PALB2, are also associated with ICLs. Moreover, we identified POLI, which is among the main polymerases involved in translesion DNA synthesis (TLS), and TFPT and CETN2, which are linked to the transcription-coupled nucleotide excision repair (TC-NER) and nucleotide excision repair (NER) pathways (Supplementary Figure 4E) [26]. Subsequently, we focused on these core DNA repair genes (GO:0006281) to gain more insights into the repair of Cas12a breaks in CAG repeats.

**Figure 2.**
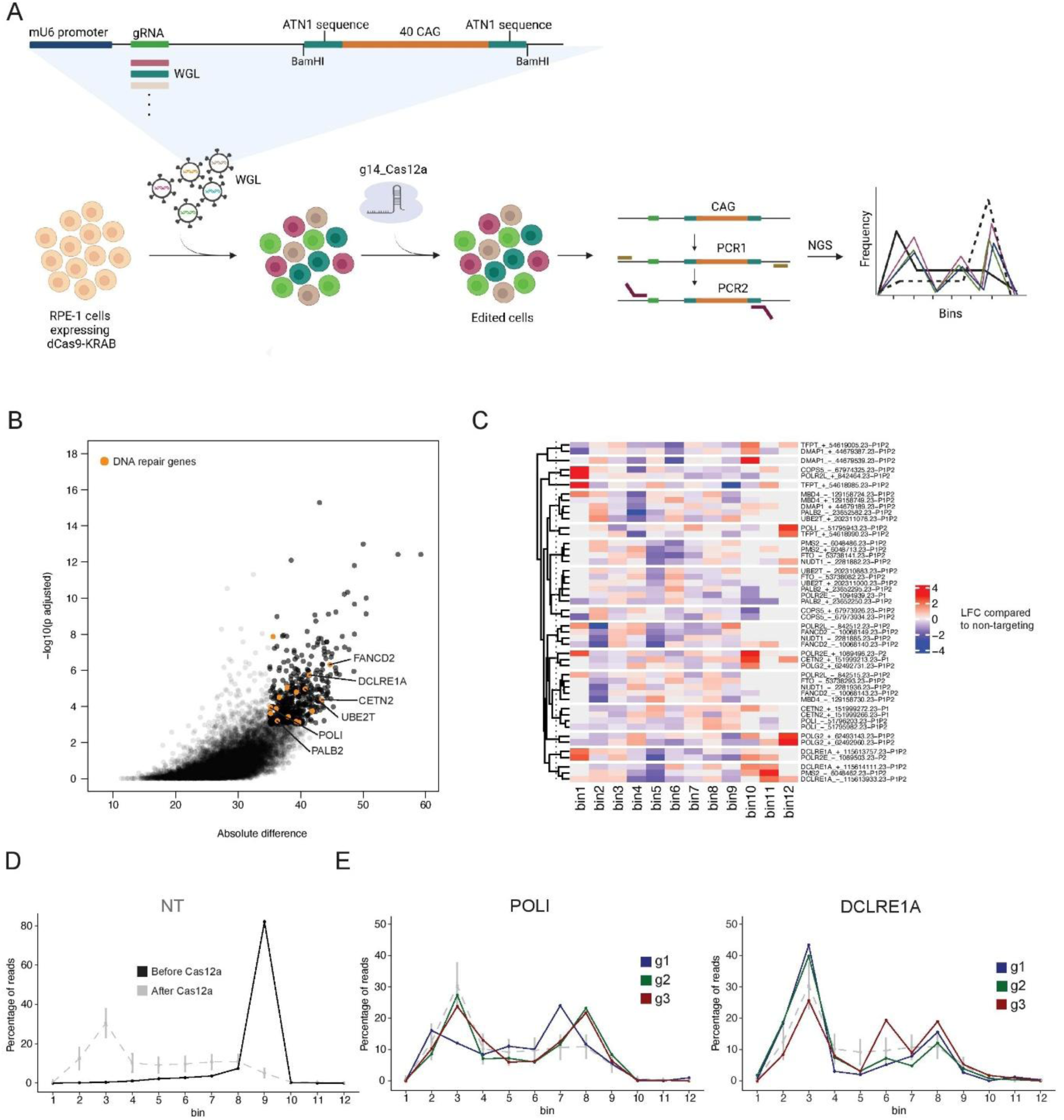
Design and outcome of the CRISPRi screen of CAG repeats. **A.** Schematic visualization of the CRISPRi screen. The Weissman CRISPRi lentiviral library (whole-genome library (WGL)) was utilized to identify genes involved in the repair of CAG repeats. gRNAs from the WGL were introduced into a vector containing the target, 40 CAG repeats, with sequences flanking the repetitive tract of the ATN1 gene. Each cell expressed dCas9-KRAB, and the library was transduced at a low MOI, allowing the integration of one gRNA-containing construct per cell. This approach facilitated the efficient inhibition of the transcription of specific genes from the library. The infected RPE-1 cell pool was nucleofected with Cas12a_g14a, and paired-end NGS of the amplified fragment (nested PCR) was performed to determine the gRNA sequence and editing outcomes at the CAG repeats. The read length included the gRNA from the WGL and 40 CAG repeats. **B.** The analysis of the CRISPRi screen using the G-test goodness of fit revealed that 466 genes significantly altered contraction patterns following Cas12a-mediated cleavage within CAG repeats. DNA repair genes are highlighted in orange. **C.** Heatmap showing the log2(fold changes) (LFCs) of all DDR genes targeting gRNAs compared with the nontargeting distribution. The results were clustered based on the number of DDR genes identified (n=16). **D.** The outcome of the CRISPRi screen was analyzed by measuring the length of the CAG repeat tract in the amplified fragment. Lengths were divided into 12 bins, resulting in a distribution of CAG repeat lengths for each gRNA. Frequency plots showing the sequence length distribution for all nontargeting gRNAs. After Cas12a cleavage, the CAG repeat contraction pattern for nontargeting gRNAs occurred, whereas before Cas12a cleavage, it corresponded to the nonedited PCR product. **E.** Frequency plots for POLI-targeting gRNAs and DCLRE1A-targeting gRNAs. The sequence length distribution is shown for three gRNAs (g1, g2, and g3).

We calculated the log2 fold change from the nontargeting distribution to confirm that gRNAs targeting the same DNA repair gene changed the contraction pattern in a similar manner. gRNAs against the same gene tended to cluster closely together (Figure 2C) when the change in distribution was used for clustering. Most of the time, two out of three gRNAs targeting genes exhibited a very similar pattern that differed from nontargeting gRNAs (Figure 2D and 2E).

We inhibited the transcription of selected genes (PALB2, UBE2T, FANCD2, CETN2, DCLRE1A, and POLI) to validate the results of the CRISPRi screen and determine which proteins may participate in the repair of Cas12a-induced DNA breaks at various stages. We analyzed the CAG contraction patterns over time via capillary electrophoresis (Supplementary Figure 6A). The amplified fragment containing 40 CAG repeats was categorized into three contraction ranges: short (40–27 CAG repeats), medium (26–11 CAG repeats), and long (10–0 CAG repeats) (Supplementary Figure 6B). In all the cases, we confirmed that the inhibition of selected candidates influenced the contraction patterns. As expected, short repeat contractions were observed at early time points and became more pronounced over time (Supplementary Figure 7). POLI and PALB2 knockdown had the most pronounced effect on the CAG contraction pattern during early editing (Supplementary Figure 6C). Interestingly, particularly at earlier time points after POLI knockdown, gradual shortening was absent, and its contraction pattern resembled that of Cas9_g10 (Supplementary Figure 6D and 7). Medium and long contractions dominated at later time points (12 h–24 h), with the highest prevalence observed in cells with FANCD2 and DCLRE1A knockdown (Supplementary Figure 6D). Even later, short contractions were more frequent in cells with the knockdown of the selected genes than in those treated with the nontargeting gRNA, confirming their involvement in the gradual shortening of CAG repeats.

### DCLRE1A acts as an endo- and exonuclease at Cas12a-generated structures within CAG repeats

Based on the results of the CRISPRi screen, we aimed to identify a factor that acts in the first stages of DNA repair, i.e., a factor that is capable of directly binding and processing structures formed by CAG repeats after DSB induction. Therefore, we selected DCLRE1A, a structure-specific 5’-3’ exonuclease that plays a crucial role in ICL repair, DSB repair, and break-induced replication at telomeres [24], [27]. In vitro, DCLRE1A also exhibits endonucleolytic activity toward single-stranded DNA [28][29]. We first investigated whether DCLRE1A accumulates at the cut site using immunocytochemistry and confocal microscopy. We confirmed the colocalization of DCLRE1A with γ-H2AX, which marks the site of the DSB (Figure 3A). A statistically significant increase of 32.91% in the colocalization rate was observed at 12 h after Cas12a_g14 electroporation compared with 13.32% in cells without DNA break induction (Figure 3A, Supplementary Figure 8). Furthermore, using ChIP‒qPCR, we detected high enrichment of DCLRE1A in the region flanking the CAG repeats following DNA break induction, confirming the direct presence of DCLRE1A near the cleavage site (Supplementary Figure 9).

**Figure 3.**
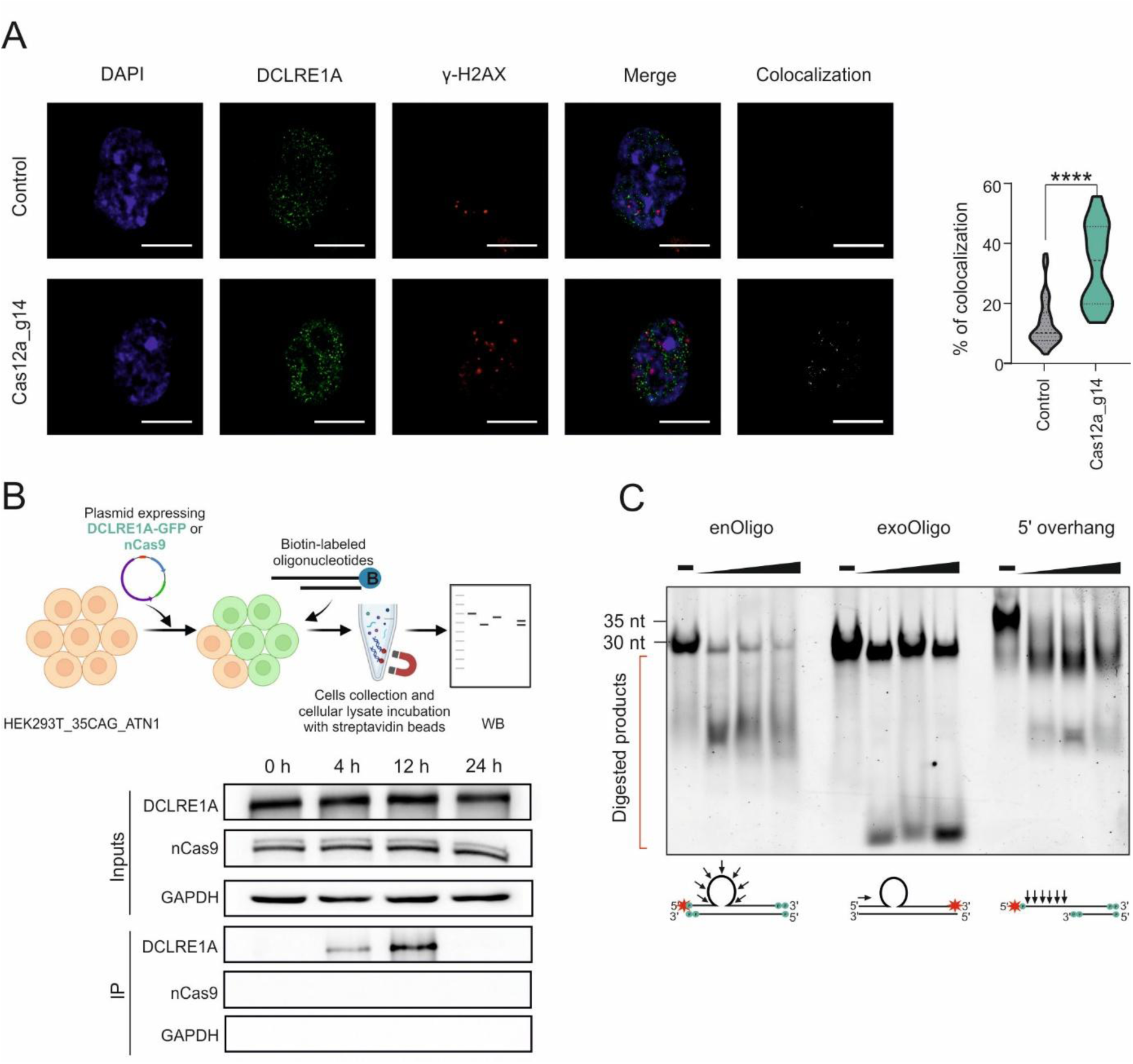
DCLRE1A recognizes and processes structures arising from 5’ overhangs. **A.** DCLRE1A accumulates at Cas12a-induced DNA breaks within a repetitive target. Representative images show the nuclei of control RPE-1 (without DSB induction) (Control) and Cas12_g14-treated RPE-1 cells infected with multiple viruses carrying 40 CAG repeats. Scale bar = 10 μm. The graph shows that the percentage colocalization coefficient increased from 13.32% to 32.91% for DCLRE1A–γH2AX colocalization following DSB induction (n = 14 and 13, respectively). For each cell, 10 to 16 micrographs were captured along the Z-axis, from which 3 micrographs were selected for calculations. Statistical analysis was conducted using the Mann‒Whitney and Kolmogorov‒Smirnov tests. Asterisks indicate levels of statistical significance: **** p ≤ 0.0001. **B.** DCLRE1A recognizes and binds to 5’ overhangs in human cells. Schematic representation of the coimmunoprecipitation experiment. DCLRE1A bound to the annealed oligonucleotides that mimic the Cas12a-generated 5’ overhangs. HEK293T_35CAG_ATN1 cells were transfected with plasmids expressing either DCLRE1A or the nickase Cas9 (nCas9) with a gRNA targeting the fragment of the *HTT* gene upstream of the CAG repeat tract (negative control). Cells were collected at 0 h, 4 h, 12 h, and 24 h following oligonucleotide delivery. **C.** The purified recombinant DCLRE1A protein exhibited endo- and exonucleolytic activities toward structures formed by 5’ overhangs in vitro. The DCLRE1A protein (25, 50, and 100 nM) was added to the annealed oligonucleotides (enOligo and exoOligo) and a positive control with a 5’ overhang. The digested products were separated on a PAGE gel under denaturing conditions. The structures arising from annealed oligonucleotides are shown below the denaturing gel. Asterisks indicate 5′-FAM-labeled nucleotides. P denotes a phosphorothioate substitution that prevents exonuclease cleavage.

We transfected HEK293T_35CAG_ATN1 cells with plasmids expressing DCLRE1A or a nickase Cas9 (negative control) and a DNA duplex resembling the structure of a Cas12a break to determine whether DCLRE1A physically binds the structures generated by Cas12a_g14 within the CAG repeats. One of the oligonucleotides from the DNA duplex was labeled with biotin, allowing the immunoprecipitation of annealed oligonucleotides with streptavidin beads (Figure 3B). This experiment showed that DCLRE1A recognized and directly bound to the oligonucleotide 4 h after its delivery into cells (Figure 3B). Interestingly, this effect was more pronounced at 12 h and was not detectable at 24 h after delivery. Therefore, we decided to explore the type of activity that DCLRE1A exerted on these structures. For this purpose, we performed an in vitro nuclease activity assay using recombinant wtDCLRE1A protein (residues 698-1040) and mutDCLRE1A (D745A, H746A) (Supplementary Figure 10) and FAM-labeled oligonucleotides (enOligo and exoOligo) that resemble the structures generated by Cas12a_g14. A DNA duplex with a T-rich 5’ overhang was used as a positive control from [28](Figure 3C). A phosphorothioate substitution at enOligo was required to prevent 5’-3’ exonuclease cleavage and analyze the endonucleolytic activity of DCLRE1A. Following annealing, enOligo and exoOligo were incubated with recombinant wtDCLRE1A or mutDCLRE1A proteins. Unlike mutDCLRE1A, wtDCLRE1A led to a decrease in the primary substrate, as reflected by the presence of digestion products on the denaturing gel (Figure 3C) (Supplementary Figure 11). The digestion pattern for enOligo suggested that oligonucleotides mimicking Cas12a-generated overhangs within CAG repeats can form a heterologous loop that might be recognized and cut by the DCLRE1A protein. In the case of exoOligo, we observed a typical pattern characteristic of the exonucleolytic activity of DCLRE1A. Thus, we showed that DCLRE1A can participate in the repair of Cas12a-induced DNA breaks within CAG repeats by binding to the structures formed after cleavage and their processing via both endo- and exonucleolytic cutting.

### DCLRE1A and POLI participate in microhomology-mediated template switching to generate inverted repeats at the cleavage site

DCLRE1A influences DNA repair by directly processing secondary structures or interacting with other proteins, such as those involved in ICLs [28], [30]. These activities of DCLRE1A are determined by its functional domains. It contains conserved catalytic MBL (metallo-β-lactamase) and β-CASP (metallo-β-lactamase-associated CPSF, Artemis, SNM1/PSO2) domains, which determine its nucleolytic activity, as well as the UBZ domain that interacts with ub-PCNA to recruit DCLRE1A to the cleavage site (Figure 4A) [31]. Since POLI knockdown inhibited the gradual shortening of CAG repeats, we investigated whether DCLRE1A and POLI are functionally connected in this process. Therefore, we performed co-immunoprecipitation (co-IP) experiments using FLAG-tagged variants of wild-type DCLRE1A (wtDCLRE1A), a catalytically inactive mutant carrying a D736A substitution within the cleavage domain (mutDCLRE1A), and a mutant lacking the UBZ domain (ΔUBZ-DCLRE1A) (Fig. 4B) to analyze their interactions with POLI following DSB induction at the CAG repeat tract. All DCLRE1A variants were found to associate with POLI, although the interaction with the ΔUBZ-DCLRE1A was reduced (Figure 4B). Moreover, consistent with the results reported by Zhang J et al., we showed that ub-PCNA immunoprecipitated with both wtDCLRE1A and mutDCLRE1A but not with ΔUBZ-DCLRE1A [27]. These findings suggest that the weaker association observed between ΔUBZ-DCLRE1A and POLI most likely results from impaired ub–PCNA–dependent recruitment of DCLRE1A to DNA damage sites. The persistence of a residual interaction signal suggests that DCLRE1A may also be capable of recognizing DNA lesions independently of the PCNA-mediated pathway. DNase treatment abolished the DCLRE1A–POLI interaction, demonstrating that the interaction is DNA-dependent (Figure 4C). Because DCLRE1A has been implicated in ICL processing, we next examined whether ICL induction influences its association with POLI. Exposure to nitrogen mustard, which increases ICL formation, enhanced the interaction between proteins as detected by co-immunoprecipitation (Supplementary Figure 12).

**Figure 4.**
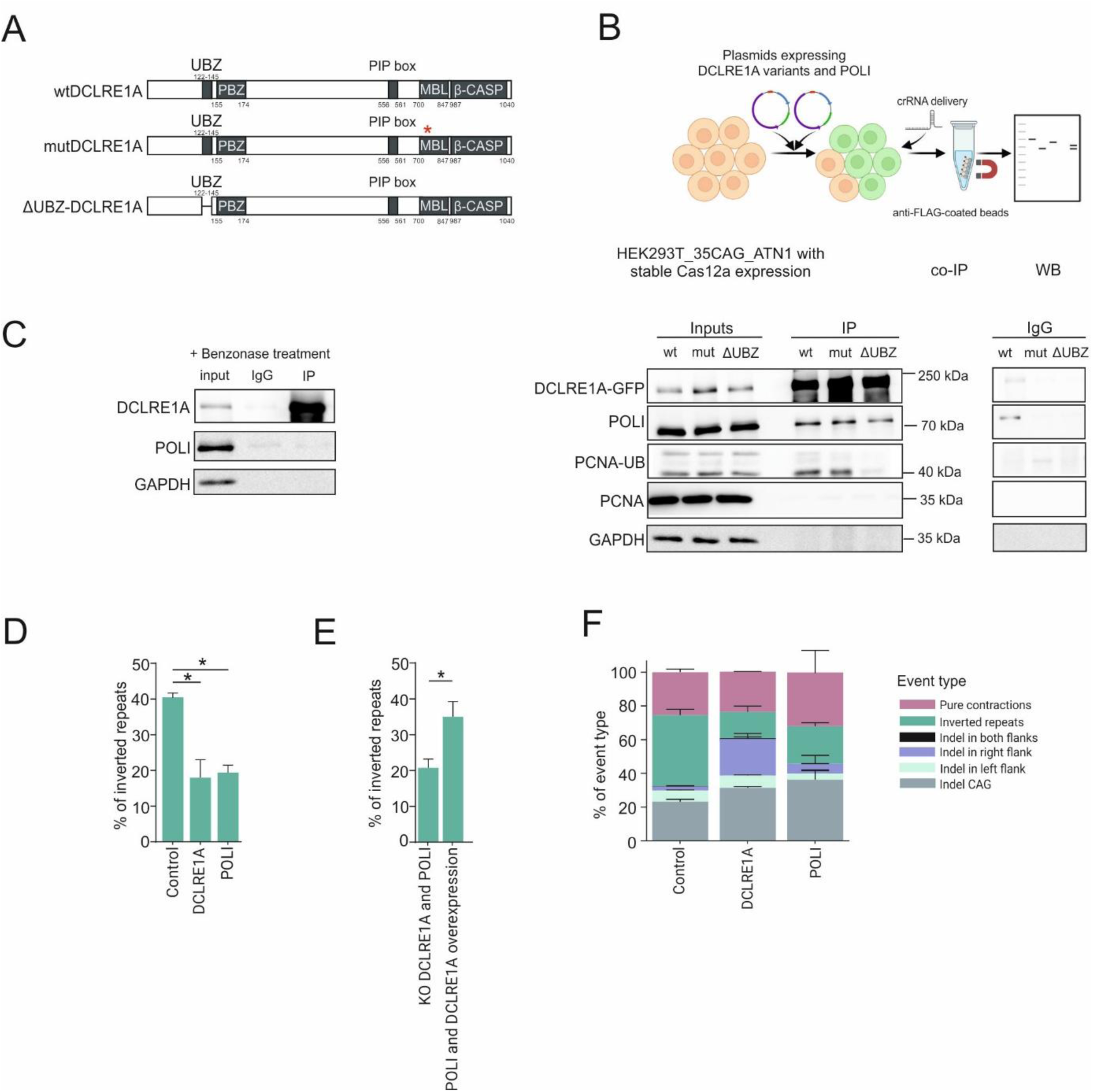
DCLRE1A-POLI influences the formation of inverted repeats. **A.** Schematic of full-length DCLRE1A and its mutants. **B.** Schematic representation of the coimmunoprecipitation procedure and coimmunoprecipitation of wtDCLRE1A, mutDCLRE1A, and ΔUBZ-DCLRE1A with POLI, ub-PCNA, PCNA, and GAPDH. **C.** Coimmunoprecipitation of Flag-tagged wtDCLRE1A with POLI following Benzonase treatment. **D.** Frequency of inverted repeats following DCLRE1A or POLI knockdown in RPE-1_40CAG cells. **E.** The results from the rescue experiments conducted in the RPEKoD and I cell lines. **F.** Analysis of the editing outcomes (NGS reads) after DCLRE1A or POLI knockdown compared with the control (nontargeting gRNA).

POLI is an error-prone polymerase that belongs to the class of TLS polymerases and rapidly repairs DSBs when the standard high-fidelity DNA polymerases cannot proceed [32]. The TLS polymerases Polζ and REV1 can synthesize perfect inverted repeats during template switch-mediated strand invasion following the generation of DNA breaks in yeast [33]. Additionally, DCLRE1A initiates end resection for template-switch-mediated strand invasion during replication, thereby maintaining the alternative lengthening of telomeres (ALT) [27]. Interestingly, after Cas12a_g14-generated DNA breaks in RPE-1_40CAG cells, nearly 40% of the NGS reads contained a mixture of inverted repeats (Figure 4D). We analyzed editing outcomes in RPE-1_40CAG cells following DCLRE1A or POLI knockdown via CRISPRi to investigate whether the POLI and DCLRE1A proteins are responsible for the introduction of inverted repeats within the CAG repeats. The analysis of the editing results revealed a statistically significant reduction of approximately 50% in inverted repeats following the knockdown of DCLRE1A or POLI compared with cells treated with a nontargeting gRNA (Figure 4D). Moreover, we conducted rescue experiments by introducing plasmids expressing DCLRE1A, POLI, or both DCLRE1A and POLI into DCLRE1A (RPE1koDCLRE1A), POLI (RPEkoPOLI), or both DCLRE1A and POLI (RPEKoD and I) CRISPR-Cas9-mediated knockout RPE-1_40CAG cell lines (Supplementary Figure 13). A statistically significant increase in inverted repeats, from 20% to 35%, was observed in cells with the simultaneous overexpression of POLI and DCLRE1A (Figure 4E, Supplementary Figure 14). These experiments supported our hypothesis that POLI-DCLRE1A is involved in the formation of inverted repeats during the template switching mechanism (Supplementary Figure 15).

Interestingly, DCLER1A knockdown did not significantly affect the frequency of pure CAG repeat contractions. However, it increased the extension of the deletion to the region flanking the CAG repeats, indicating its role in maintaining the deletion within the CAG repeat tract (Figure 4F). This result implies that, in addition to its role in template switching mechanisms, DCLRE1A may participate in other DNA repair pathways, including those leading to pure short deletions within the repetitive tract.

### DCLRE1A cooperates with SLX4 during Cas12a-generated DNA break repair to produce pure CAG repeat contractions

We assessed the final truncation products, i.e., variants with 2 and 3 CAG repeats, which constituted ∼40% of all pure contracted variants in RPE-1_40CAG cells following Cas12a-mediated DNA breaks, to better understand the mechanisms leading to the formation of pure repeat contractions. We filtered genes from the CRISPRi screen for which knockdown reduced the occurrence of these contractions to less than 5% to identify genes involved in the generation of these variants. Interestingly, among the filtered genes, we identified DNA repair genes related to the regulation of DNA replication, namely, CCNA2, INO80B, MCM3, RAD17, and ZNF830, and genes associated with FA and ICL repair, namely, FANCG, FANCB, and SLX4, indicating their potential roles in the pure gradual shortening of CAG repeats (Figure 5A). SLX4 acts as a molecular scaffold that interacts with the structure-specific endonucleases XPF-ERCC1, MUS81-EME1, and SLX1 through its MLR, SAP, and CCD domains, respectively [34], [35]. These complexes cleave a wide range of branched DNA structures produced during the repair of damaged DNA and broken DNA replication forks, such as D-loops formed after strand invasion and DNA repair and Holliday junctions (HJs) during HR. Because DCLRE1A and SLX4 are thought to cooperate in the processing of DNA repair intermediates, we investigated how this cooperation influenced the generation of pure contracted CAG repeat variants. We found that SLX4 knockdown alone and the simultaneous knockdown of SLX4 and DCLRE1A caused statistically significant decreases in those contracted events (∼50%), with even greater statistical significance observed after the simultaneous knockdown of both genes (Figure 5B, Supplementary Figure 16).

**Figure 5.**
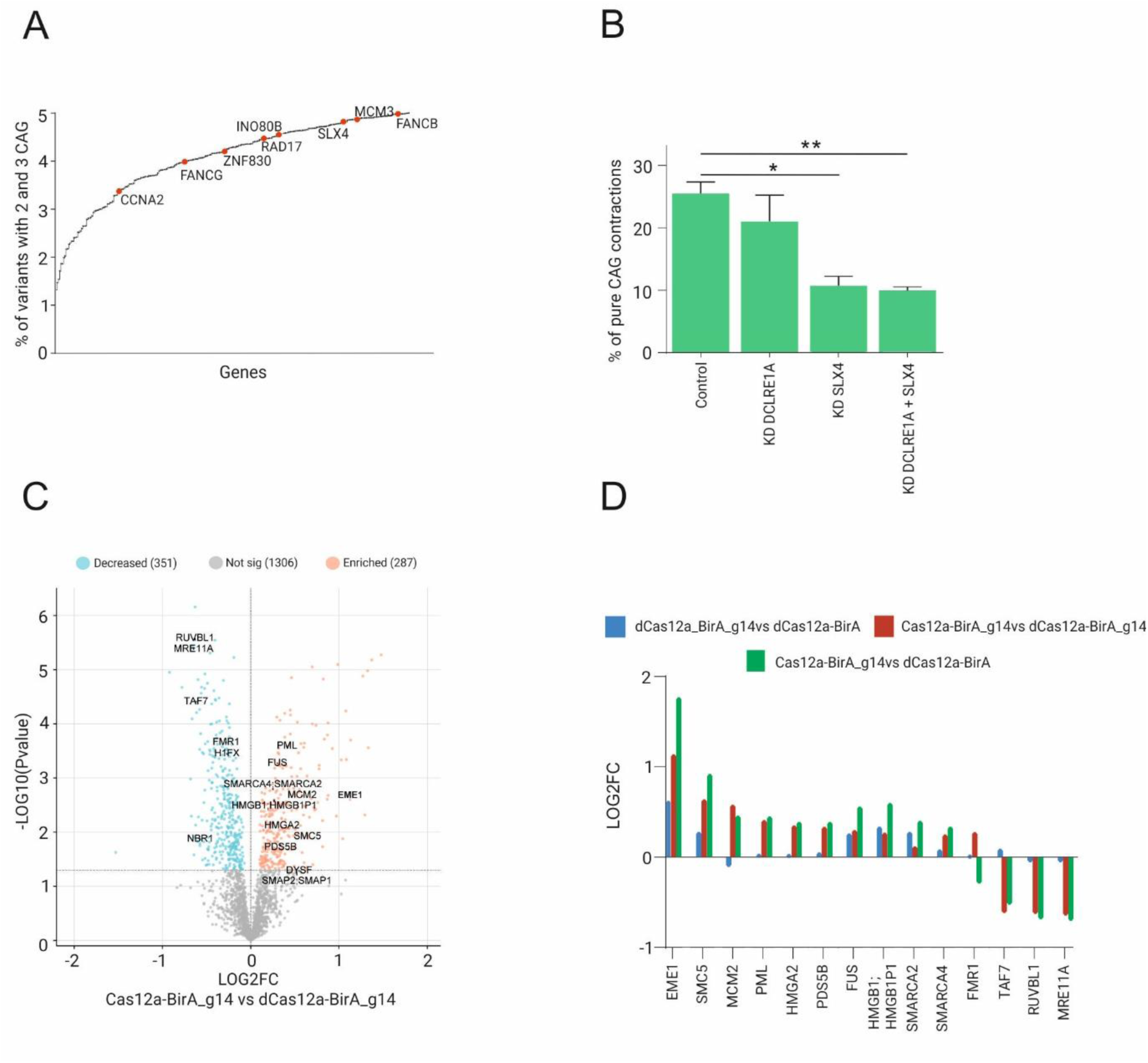
SLX4 and DCLRE1A influence the occurrence of pure contraction events following Cas12a-induced DNA breaks. **A.** The graph shows DNA-related genes whose knockdown resulted in a drastic decrease in the number of events with 2 and 3 CAG repeats. **B.** The graph shows the percentage of pure contractions following the knockdown of DCLRE1A (DCLRE1A), SLX4 (SLX4), and both DCLRE1A and SLX4 (DCLRE1A + SLX4). The bars in the graph represent the mean values of the pure contracted variants ± SEMs. The p-values are indicated by asterisks (*p < 0.05 and **p < 0.01). This experiment was repeated twice. **C.** Volcano plot showing DNA repair-related genes detected in the BioID analysis whose expression decreased or was enriched after Cas12a-induced DNA breaks within CAG repeats. Here, we compared cells expressing Cas12a (Cas12a-BirA_g14) and cells expressing dCas12a (dCas12a-BirA_g14), both of which were treated with g14. **D.** Comparison of the cleavage effect(Cas12a_BirA_g14 vs. dCas12a_BirA_g14), and binding effect (dCas12a_BirA_g14 vs. dCas12a_BirA) for only the DNA-related proteins identified in the BioID analysis.

In addition, using proximity-based proteomics (BioID and BirA*), we confirmed the presence of the SLX4-interacting protein EME1 near the Cas12a_g14 cutoff site (Figure 5C and 5D; Supplementary Figure 17). Moreover, we also observed the statistically significant enrichment (q value <0.05) of the SMC5, PML, HMGA2, PDS5B, SMARCA2, SMARCA4, and FUS proteins related to HR, where FUS is engaged in D-loop formation, and PDS5B can preferentially bind with strong affinity to such structures [36], [37]. The enrichment of those proteins was directly related to Cas12a-mediated DSBs; only a slight increase was detected in dCas12a-BIRA_g14 cells (Figure 5D). Interestingly, in a BioID analysis (4 biological replicates), a decrease in the level of MRE11, the key factor responsible for the end resection required for classic HR, was observed [38]. The presence of SLX4-interacting proteins alongside other HR-associated factors suggests that SLX4 and DCLRE1A participate in CAG repeat shortening, resulting in pure contractions.

### Controlled contraction of mutant CAG repeats using Cas12a fused to DNA repair factors

Depending on the cell type, pure contracted variants account for 20% to 50% of the overall events after Cas12a-mediated editing. Our research revealed proteins that influenced the formation of pure contracted variants. Therefore, we wondered whether their presence at the cleavage site could increase the number of pure CAG deletions in the endogenous locus. We fused Cas12a with FANCG, PALB2, and DCLRE1A, and the SLX4 domains, SAP and MLR. This approach ensures the presence of DNA repair factors directly at the cleavage site, thereby enhancing their direct participation in repairing Cas12a-induced DNA breaks within the repetitive tract (Figure 6A). Plasmids expressing the fusion proteins were introduced into HEK293T_35CAG_ATN1 cells, and the editing process was initiated following g14 delivery (Figure 6A). According to the NGS data, we observed approximately 90% editing efficiency after all the fused proteins were used, which was comparable to that of Cas12a alone, suggesting that they did not influence the editing process. Interestingly, FANCG, SAP, MLR, and PALB2 increased the occurrence of pure shortened variants after cutting (Figure 6B). The greatest increase, approximately 16%, was observed for Cas12a-MLR (Figure 6B). Moreover, Cas12a-FANCG and Cas12a-MLR fusions decreased the extension of the deletions to the site flanking the CAG repeats (Figure 6B). Only 48% pure contraction events were observed following Cas12-DCLRE1A-mediated cleavage because of the increase in the number of inverted repeats at the break site (Supplementary Figure 18).

**Figure 6.**
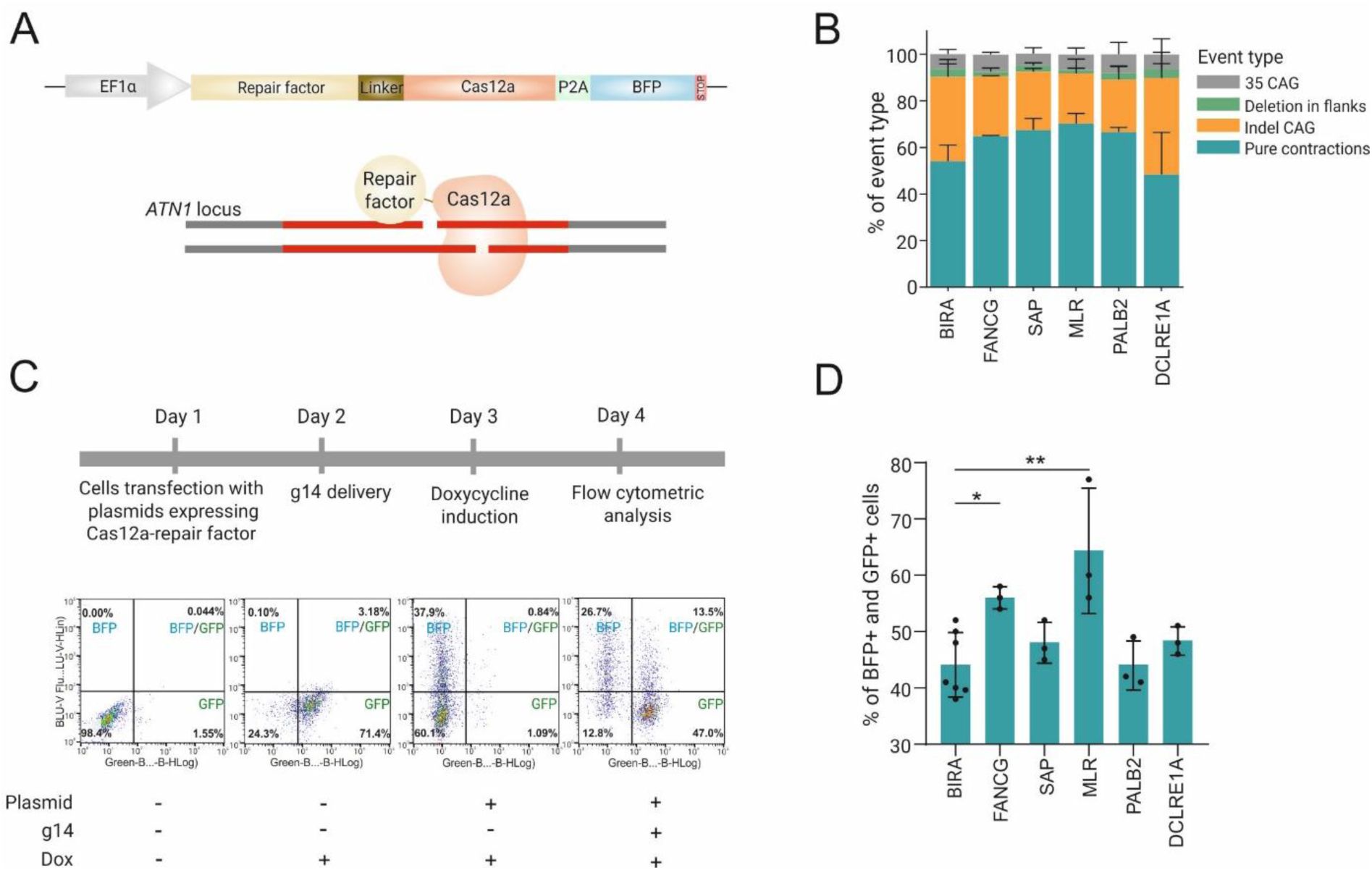
Fusion proteins increase the number of pure contractions. **A.** Schematic representation of the vector expressing the Cas12a-DNA repair factor fusion protein. **B.** NGS data revealed variations in the contribution of different types of events mediated by the Cas12a-DNA repair factor fusion protein in HEK293T_35CAG_ATN1 cells. BIRA-treated cells served as controls. **C.** Timeline of the experiment for determining the frequency of pure contraction events after Cas12a-DNA repair factor treatment in HEK-Flp-In_T-Rex_64CAG cells. Representative flow cytometry plots are shown below. The first plot displays untreated cells, whereas the second shows cells treated with doxycycline. The third and fourth plots show cells treated with plasmids expressing the fusion protein and BFP and cells treated with plasmids and g14, respectively. **D.** The graph shows the percentage of GFP-positive cells (pure contraction variants) following treatment with the fusion proteins. The statistical analysis was performed using an unpaired t-test (n = 3), with a p-value of ≤ 0.0001.

Since our strategy was verified using a model with 35 CAG repeats, we investigated the contraction events associated with 64 CAG repeats, which are considered pathological in nearly all polyQ diseases. We created a HEK293 cell line in the Flp-In T-REx system with doxycycline-inducible expression of a fragment of the *ATN1* gene containing 64 CAG repeats fused to the GFP sequence (HEK-Flp-In_T-Rex_64CAG). If the number of CAG repeats was shortened out-of-frame during DNA repair, the number of GFP/BFP-positive cells following doxycycline induction would decrease. Therefore, using flow cytometry, we calculated the number of pure shortened variants as a population of BFP/GFP-positive cells compared with BFP-positive cells (plasmid expressing Cas12a-repair factor) after DNA break induction (Figure 6C). We introduced plasmids expressing Cas12a fused with repair factors into HEK293-Flp-In_T-Rex_64CAG cells and induced DSBs following g14 delivery (Figure 6D). Interestingly, in this assay, we also observed statistically significant increases in pure shortened variants of 16% and 20% for Cas12a-FANCG and Cas12a-MLR, respectively. Cas12a-DCLRE1A and Cas12a-SAP exhibited slight, non-significant increases, and the BFP/GFP signals of Cas12a-PALB2 were the same as those of the control (Figure 6D). These findings indicated that the Cas12a-MLR fusion can be successfully used to shorten CAG repeats, offering a new perspective on targeting repetitive sequences.

## DISCUSSION

The expansion of short tandem repeats underlies many currently incurable human diseases. Controlled shortening of these sequences to nonpathological lengths would eliminate the direct cause of the disease and provide a universal therapeutic approach. Here, we show for the first time that Cas12a can be used for this purpose and that it represents a promising alternative to Cas9. In our model system utilizing CAG repeats in the context of the *ATN1* gene, AsCas12a caused a gradual shortening of the repeats. Interestingly, in a previous study investigating endonuclease efficiency in targeting various tandem repeats in a yeast model, *Francisella novicida* Cas12 (FnCas12a) showed no activity against CAG/CTG repeats, and only GAA triplets were efficiently edited [39]. Our detailed analysis of Cas12a editing outcomes revealed two types of products, namely, pure contractions and contractions accompanied by the insertion of inverted repeats, the number of which increased with time. Because these insertions can result in gene knockout, their presence is undesirable from a therapeutic standpoint. We reasoned that mechanistic insights into the repair mechanism not only could allow us to control the editing outcomes but will also be essential for the translation of this approach into the clinic.

Previous research has suggested that Cas9-induced DSBs within CAG/CTG repeats are repaired mainly by NHEJ, HDR, and SSA [8][10]. NHEJ also predominates during the repair of nonrepetitive sequences edited by Cas12a [40]. The mechanisms responsible for the repair of staggered cuts generated by Cas12a directly within the MH region are unknown. Therefore, we developed a CRISPRi screen to improve our understanding of DSB repair at MH sequences, enabling the first unbiased, genome-wide identification of DNA repair factors involved in their repair. Notably, this method can be adapted to study CAG repeat instability induced by different endonucleases.

Surprisingly, a CRISPRi screen revealed a central role for ICL repair factors in mediating Cas12a-induced CAG repeat contraction, with DCLRE1A emerging as a key effector. Mechanistically, our in vitro data support a model in which the 5′ overhang generated by Cas12a cleavage forms a heterologous loop recognized by DCLRE1A, initiating the deletion of one or several CAG units. We further demonstrated that DCLRE1A binds directly to CAG-rich structures in human cells and interacts with additional repair proteins identified in our CRISPRi screen, including POLI and SLX4. According to previous studies, FANCD2 and PALB2, which were also identified in our screen, may act in the same repair pathways as DCLRE1A. Since UBE2T-mediated ubiquitination promotes the recruitment of the FANCD2–FANCI complex to sites of replication stress, FANCD2 cooperates with PALB2 and SLX4 in downstream steps of ICL repair through HR [26], [41], [42]. In our context, the source of replication stress may be secondary structures formed by expanded CAG repeats or the presence of Cas12a bound to DNA, both of which could serve as signals for the activation of the ICL repair machinery.

Moreover, we found that DCLRE1A knockdown reduced but did not completely halt gradual shortening. Importantly, its inhibition caused a significant increase in deletions extending to the site flanking the CAG tract, indicating that DCLRE1A is crucial for maintaining short deletions within the repeats. In contrast, POLI knockdown completely inhibited gradual shortening, suggesting that it affects both pure contracted variants and those with inverted repeats. Since POLI lacks exo- and endonuclease activities, its role in the mechanism of repeat contraction remains elusive and requires further investigation. Current knowledge of the participation of POLI in DNA repair is limited. It belongs to the class of TLS polymerases, which are recruited to bypass damage and continue DNA synthesis in regions that are difficult to replicate, such as those containing bulky adducts, secondary structures, or DNA–protein cross-links [43], [44]. We are the first to report the cooperation of POLI with DCLRE1A and its direct involvement in the repair of DSBs within repeated sequences. A rescue experiment revealed that the interaction between POLI and DCLRE1A is crucial for the formation of inverted repeats. In addition to POLI, DCLRE1A also precipitated with the scaffold protein SLX4, and inhibiting both proteins simultaneously significantly reduced the number of pure contracted variants. The SLX4-interacting protein EME1 and other HR-related proteins were detected by BioID analysis, indicating a mechanistic model in which pure contracted variants arise through a coordinated HR-based pathway involving POLI, DCLRE1A, and SLX4. Notably, the fusion proteins we developed, especially Cas12a-MLR, significantly increased the number of pure contractions for both 40 and 64 CAG repeats. In addition, Cas12a-MLR markedly reduced the occurrence of deletions in the sequences flanking the CAG tract. The relatively small size of the MLR domain (6.44 kDa) facilitates the packaging of the Cas12a-MLR construct into viral vectors, a highly desirable feature for potential therapeutic applications. Our data also indicate that the duration of Cas12a-mediated editing influences the frequency of pure contracted variants and the length of the CAG tract. Therefore, an alternative method to increase the proportion of pure contractions could be to stop the editing process in its early stages.

Among the limitations of this study is the design of the CRISPRi screen, which did not permit the analysis of larger deletions. In addition, future work should investigate this approach in postmitotic neurons, where repeat instability contributes to neurodegeneration. Overall, our data indicate that the Cas12a-induced DSBs within repeated sequences engage noncanonical DNA repair mechanisms that differ from those operating in nonrepeating sequences. We have shown that this knowledge can be used in practice to increase the number of desired editing outcomes.

## MATERIAL AND METHODS

### Cell culture

Human embryonic kidney cells (HEK293T) and HEK-Flp-In_T-Rex_64CAG cells were cultured in Dulbecco’s modified Eagle’s medium (Lonza) supplemented with 10% fetal bovine serum (Biowest), antibiotic–antimycotic solution (Sigma‒Aldrich), and 2 mM L-glutamine (Sigma‒Aldrich). Additionally, the medium of Flp-In T-REx-293 cells was supplemented with 100 µg/ml hygromycin B (Thermo Fisher Scientific) and 5 µg/ml blasticidin S (Thermo Fisher Scientific). hTERT-immortalized retinal pigment epithelial cells (hTERT RPE-1) were cultured in Dulbecco’s modified Eagle’s medium/nutrient mixture F12 (DMEM-F12) (Thermo Fisher Scientific) supplemented with 10% fetal bovine serum (Sigma‒Aldrich) and antibiotic–antimycotic solution (Sigma‒Aldrich).

### Generation of HEK293T_35CAG_ATN1 cells

The donor template for HDR was a linearized plasmid, pUC57, with 35 CAG repeats and sequences flanking the CAG repeats from the *ATN1* gene (411 bp and 395 bp from the repetitive tract; listed in Supplementary Material 1), cloned at the EcoRI and HindIII restriction sites. A silent mutation was added 3 nt downstream of the PAM sequence in the donor template (GTC to GTT) to prevent re-cleavage. Three micrograms of the plasmid expressing sgATN1 and 2 μg of the linearized donor template were electroporated into HEK293T cells using the Neon™ Transfection System (Invitrogen) in 100 μl tips with the following parameters: 1150 V, 20 ms, and two pulses. Cells were sorted based on the GFP signal using a FACSAria device (Becton Dickinson) into 96-well plates (one cell per well) and cultured until they reached 50% confluence. Next, each clone was harvested, and the CAG repeat length was checked by amplifying the fragment containing the repetitive tract using ATN1_F and ATN1_R primers (listed in Supplementary Table 2). Protein expression was evaluated by Western blot analysis with an anti-ATN1 antibody.

### Electroporation of RNP complexes

Cas9_g10 and Cas9_g11 complexes were prepared according to the instructions of the IDT Alt-R CRISPR-Cas9 System. Briefly, the crRNA and tracrRNA oligos were mixed in equimolar concentrations to a final duplex concentration of 44 μM (g10, g11). The mixture was heated at 95 °C for 5 minutes and incubated at room temperature for 20 minutes. Then, the guide complex was mixed with the SpCas9 enzyme (VBCF Protein Technologies facility http://www.vbcf.ac.at) at a ratio of 22 pmol to 18 pmol. The mixture was incubated at room temperature for 20 minutes.

Cas12a_g14 was formed by mixing 6.33 Alt-R™ A.s. Cas12a (Cpf1) Ultra (IDT) with 20 pmol of g14 and incubating the sample for 20 min at room temperature. A total of 20 pmol of Alt-R Cas12a (Cpf1) electroporation enhancer (IDT) was added to RNP complexes before electroporation to improve the efficacy of electroporation. HEK293T_35CAG_ATN1 and hTERT RPE-1_40CAG cells were electroporated using the Neon™ Transfection System (Invitrogen). Briefly, 2.5 × 10^5^ HEK293T_35CAG_ATN1 cells or hTERT RPE-1_40CAG cells were harvested, resuspended in PBS, and electroporated with RNP complexes in 100 μl tips using the following parameters: 1150 V, 20 ms, 2 pulses or 1650 V, 10 ms, and 3 pulses for HEK293T_35CAG_ATN1 and hTERT RPE-1_40CAG cells, respectively. The crRNA sequences are listed in Supplementary Table 3.

### CRISPRi screen

A human genome-wide CRISPRi-v2 library (Addgene #83969), consisting of 104,535 gRNAs targeting the transcription start site (TSS) of ∼ 20,000 genes, was used. The target, 40 CAG repeats with 72 bp and 50 bp flanking sites, was cloned and inserted into the transfer plasmid (Addgene #84832) at the BamHI restriction site. WGL was transferred into the transfer plasmid following WGL amplification using Q5 High-Fidelity 2X Master Mix according to the manufacturer’s instructions (NEB). The PCR product was purified using a gel purification and extraction kit (Machery-Nagel, NucleoSpin Blood L or XL, depending on the cell number), and 1.9 ng of the BlpI–BstXI-digested insert was ligated to 500 ng of the digested transfer plasmid. The mixture was incubated at 16°C for 16 hours. Next, the ligation reaction was purified using ethanol precipitation. Electrocompetent MegaX DH10B T1R Electrocomp™ cells (Thermo Fisher Scientific) were utilized for large-scale transformation. A fragment of the transfer plasmid encompassing the gRNA sequence and the target was amplified using Q5 High-Fidelity 2X Master Mix (NEB) in a nested PCR approach to analyze the distribution of the gRNAs and the length of the repeats. Using primers 171 and 172, an approximately 800 bp sequence was amplified. In the second PCR, primers containing Illumina overhangs and barcodes were used to amplify an ∼350 bp product. Next, the PCR product was purified using solid-phase reversible immobilization (SPRI) beads and subjected to Illumina sequencing (MiSeq, 600 cycles).

For lentivirus production, the transfer plasmid was cotransfected with the packaging plasmids pCMV-VSVG (Addgene #8454) and pCMV-dR8.2 dvpr (Addgene #8455) into HEK 293T cells. Approximately 83 million RPE-1 cells were infected with the lentivirus at an MOI of 0.3, followed by approximately one week of puromycin selection (15 µg/ml). The cells were split daily until they exhibited approximately 100% BFP signal. Twenty-two million cells were electroporated with Alt-R™ A.s. Cas12a (Cpf1) Ultra (IDT) and g14-targeted CAG repeats using the P3 Primary Cell 4D-Nucleofector™ X Kit L (Lonza). The cells were cultured for 4 days after electroporation. The Gentra Puregene Kit (Qiagen) was used to extract DNA. DNA from cells treated with Cas12a_g14 and untreated cells was amplified using nested PCR with NEBNext^®^ Ultra™ II Q5^®^ Master Mix with 171 and 172 primers and a second PCR with MD1, MD2, MD3, MD4, and oMFM_201 primers. PCRs were pooled, purified using SPRI beads, and analyzed on a 1.5% agarose gel. After editing, NextSeq 400 M reads with a 300 bp read length were used to verify the gRNA sequences and the lengths of the CAG repeats.

### Bioinformatic analysis of the CRISPRi screen

Paired-end reads were merged into a single file using the mergepe command from seqtk (v1.3-r106) with the default arguments. For each read, forward and reverse anchors were searched with hamming distances of ≤1 and ≤2, respectively. For reads with both anchors present, we extracted the barcode, calculated the sequence length between anchors and generated a count table for all present barcodes and insert sizes. Insert sizes between the anchors were used to assess repeat lengths, and sequence lengths were sorted into bins. The bin number was optimized using the Freedman–Diaconis rule for all the NT guides and resulted in 12 bins (breaks = -1, 20, 40, 60, 80, 100, 120, 140, 160, 180, 200, 220, and 241)[45]. Replicates were combined, and the NT distribution was used as the expected distribution to compare the repeat length distribution of each targeted sgRNA using the G-test of goodness of fit [23]. The three most significantly different sgRNAs per gene were selected, and pooled G tests and G tests of heterogeneity were performed. The effect size was measured by calculating the absolute difference between each sgRNA Abs_dif gRNA 𝐴𝑏𝑠_𝑑𝑖𝑓_ (𝑠𝑔𝑅𝑁𝐴) = ∑_k=1_^12^ | 𝑑^𝑘^| and the NT guide distribution (k = bin number). The mean absolute difference was calculated as 𝑚𝑒𝑎𝑛(∑_i=1_^3^ 𝐴𝑏𝑠_𝑑𝑖𝑓_(𝑠𝑔𝑅𝑁𝐴)^𝑙^), where l is the sgRNA ID. The p values from the pooled G test were adjusted for the false discovery rate using the Benjamini‒Hochberg (FDR) procedure. The graph for Figure 2b was generated using plotly (4.10.4). The GO term GO:0006281 was used to detect genes involved in DNA repair. The log2(fold changes) of the frequencies were calculated against the NT distribution and plotted using ComplexHeatmap (2.24.0). For frequency plots, the means and standard deviations of all nontargeting sgRNAs were plotted with each separate targeting sgRNA using ggplot2 (3.5.2). Mageck was used to calculate log2(fold-change) and adjusted p values for the screen QC [46]. Essential genes were defined based on the essential gene list reported by Hart et al [47].

### Lentivirus production and cell infection

For lentivirus production, a modified version of plasmid #84832 containing a fragment of the *ATN1* gene with 40 CAG repeats at the BamHI restriction site and gRNAs from the CRISPRi-v2 Library (listed in Supplementary Table 4) was cotransfected with the packaging plasmids pCMV-VSV-G (Addgene #8454) and pCMV-dR8.2dvpr (Addgene #8455) in HEK293T cells. The medium was collected on days 2 and 3 posttransfection, and the viral supernatants were passed through 0.45-μm filters. Twenty-four hours before transduction, 6 × 10^5^ RPE-1 cells were plated in each well of a 6-well plate. RPE-1 cells were transduced at an MOI of 0.3 in the presence of polybrene (4 μg/mL). At 24 h postinfection, the medium was replaced, and at 48 h post-transduction, 15 µg/mL puromycin (InvivoGen) was added. Three days after puromycin selection, the cells were harvested for further analysis. RPE-1 cells infected with a virus carrying 40 CAG repeats are referred to as RPE-1_40CAG in this work.

### RNA extraction and RT‒qPCR

Total RNA was isolated from RPE-1_40CAG cells using TRI Reagent (BioShop) according to the manufacturer’s instructions. The RNA concentration was measured using a spectrophotometer (DeNovix). A total of 700 ng of RNA was reverse transcribed at 55 °C using Superscript III (Life Technologies) and random hexamer primers (Promega). Complementary DNA (cDNA) was employed for quantitative polymerase chain reaction (qPCR) using SsoAdvanced™ Universal SYBR^®^ Green Supermix (Bio-Rad) with denaturation at 95 °C for 30 s, followed by 40 cycles of denaturation at 95 °C for 15 s and annealing at 60 °C for 30 s in the CFX Connect™ Real-Time PCR Detection System (Bio-Rad). The sequences of the primers are presented in Supplementary Table 2. Data preprocessing and normalization were performed using Bio-Rad CFX Manager software (Bio-Rad).

### PCR product analysis by capillary electrophoresis

RPE-1_40CAG cells were electroporated with Cas12a_g14 and were collected at 0 h, 4 h, 6 h, 8 h, 12 h, and 24 h after Cas12a_g14 delivery. DNA was isolated using QuickExtract DNA Extraction Solution (Biosearch Technologies) according to the manufacturer’s instructions. NEBNext^®^ Ultra™ II Q5^®^ Master Mix with primers screen2F and screen2R was used to amplify DNA fragments containing CAG repeats under the following conditions: 98 °C for 30 s; 32 cycles of 98 °C for 7 s, 68 °C for 15 s, and 72 °C for 15 s; and 72 °C for 2 min. The CAG repeat contraction patterns were assessed by capillary gel electrophoresis (Tape Station System) using D1000 ScreenTape & Reagents (Agilent) and analyzed with TapeStation Analysis Software 4.1.1 (Agilent).

### Immunofluorescence staining and microscopy

RPE1 cells were transduced with a lentivirus carrying 40 CAG repeats (without BFP) at an MOI of 10, which increased the number of target sites, to improve visualization. A total of 5 × 10^4^ infected RPE-1_40CAG cells (without BFP) were electroporated with Cas12a_g14 and seeded on glass coverslips (Bionovo) coated with Geltrex (Thermo Fisher Scientific) in 12-well plates. After 12 hours, the cells were fixed with 4% PFA (Thermo Fisher, Paraformaldehyde, 32% w/v aqueous solution, methanol-free, Cat: 047377.9L) for 30 minutes at room temperature. The fixed samples were washed with PBS for 10 min, permeabilized with 0.5% Triton X-100 in PBS and blocked with blocking buffer (1% BSA and 0.2% Triton X-100 in PBS) for 1 h at room temperature. Cells on the coverslips were incubated overnight at 4 °C with the primary antibodies rabbit anti-DCLRE1A (Thermo Fisher Cat: PA5-57868) and mouse anti-phospho-H2A.X (Ser139) antibody (BioLegend Cat: 613401) diluted in blocking buffer. The next day, the cells on the coverslips were washed three times with PBS and incubated with the secondary antibodies anti-rabbit Alexa Fluor 488 (Jackson ImmunoResearch) and anti-mouse Alexa Fluor 594 (Jackson ImmunoResearch) for one hour at room temperature in the dark. Cells on the coverslips were washed three times with PBS and then transferred to SuperFrost Plus microscope slides (Thermo Fisher Scientific) using SlowFade Diamond Antifade Mountant with DAPI (Life Technologies). In addition to the treated samples, control samples without Cas12a_g14 treatment were prepared.

#### Settings

Micrographs were captured using a Leica TCS SP5 II confocal microscope (63x 1.4 objective, oil immersion with a detector magnification of 8, which allowed the size of each image to be set to a constant value of 30.75 µm × 30.75 µm). Laser line 488 nm, with 16% of the total laser power, and laser line 594 nm, with 25% of the total laser power, were used for the colocalization study. In this study, standard PMT detectors included in the microscope were used. All the images were captured with the same parameters during the two microscopy sessions.

#### Colocalization parameters

In this experiment, 13 study cells and 14 control cells were analyzed. For each cell, a series of 10 to 16 micrographs (depending on cell thickness) were captured along the Z axis, from which 3 micrographs were selected for each cell for the calculations. The selection of images relies on the position of the cell nucleus in the cell, and images lying next to each other on the Z axis were never selected because of the possibility of repeating the exact measurement. Colocalization was measured in all the samples using the same threshold (30%) and background (20%) parameters, which was possible using a standardized micrography procedure. The results are presented as the percentage colocalization coefficient over the entire micrograph surface.

#### Software

Calculations were performed using Leica Application Suite X version 3.4.1.18368, and statistical analyses were performed in GraphPad Prism version 9.0.

### Immunoprecipitation

A total of 5x10^6^ RPE-1_40CAG cells were electroporated with RNP complexes (Cas12a_14). Formaldehyde (Thermo Scientific) at a final concentration of 1% was used for cross-linking. The reaction was stopped by the addition of glycine (BioShop) at a final concentration of 125 nM. Cells were then collected by scraping in cold PBS supplemented with 1 mM PMSF. After centrifugation, the nuclei were isolated with cellular lysis buffer (5 mM PIPES (pH 8), 85 mM KCl, and 0.1% Triton-1000x). The nuclei were lysed in RIPA buffer (50 mM Tris-HCl (pH 8), 150 mM NaCl, 2 mM EDTA (pH 8), 1% Triton X-100, 0.5% sodium deoxycholate, and 0.2% SDS) supplemented with 1 mM PMSF. The lysates were sonicated using a BioRuptor Pico sonicator (Diagenode; 5 cycles, 30 s ON, 30 s OFF; the average length of the chromatin fragments was 400 bp). Finally, the lysates were centrifuged (12500xg, 4 °C, 5 min), and the supernatants were collected and transferred to new tubes. The lysates were suspended in RIPA buffer supplemented with 1 mM PMSF and incubated with magnetic beads (Dynabeads^TM^ Protein G, Invitrogen) coated with an antibody against DCLRE1A (listed in Supplementary Table 5) overnight at 4 °C with gentle rotation. The magnetic beads were subsequently washed 6 times with RIPA buffer and 1 time with TBS (Pierce). The beads were incubated in elution buffer (50 mM Tris-HCl (pH 8), 10 mM EDTA (pH 8), and 1% SDS) at 65 °C for 5 h to elute the immunoprecipitated proteins. The eluates were further subjected to either ChIP‒qPCR analysis or Western blot analysis.

### ChIP‒qPCR

The eluates were incubated with TE buffer and proteinase K (Diagenode) at 55 °C for 1 h. A GeneJET PCR Purification Kit (Thermo Fisher Scientific) was used to purify the DNA. The input was prepared as described above to normalize the chromatin concentration; however, the incubation with magnetic beads was omitted. Precipitated DNA and input samples were analyzed by quantitative PCR using a CFX Connect Real-Time PCR Detection System (Bio-Rad) and SsoAdvanced Universal SYBR Green Supermix (Bio-Rad). The regions downstream of the CAG repeat tract and a reference gene, β-actin, were amplified under the following thermal cycling conditions: denaturation at 95 °C for 30 s, followed by 40 cycles of denaturation at 95 °C for 15 s and annealing and elongation at 60 °C for 30 s. The primer sequences are listed in Supplementary Table 2. Input recovery was determined based on the average quantification cycle (Cq) for reactions for the precipitated DNA and input. Fold enrichment was calculated based on the ratio between input recovery for the region downstream of the repeat tract and the reference gene.

### Coimmunoprecipitation of DCLRE1A with biotin-labeled oligonucleotides

HEK293T_35CAG_ATN1 cells with stable Cas12a expression (2.5 × 10^6^) were transfected with 4 µg of a plasmid expressing FLAG-tagged DCLRE1A using PEI. Forty-eight hours after transfection, 6 × 10^6^ cells were electroporated with 5 nmol of annealed oligonucleotides using the Neon transfection system (the sequences of the oligonucleotides are listed in Supplementary Table 6). Briefly, cells were resuspended in PBS and mixed with annealed oligonucleotides. Electroporation was then performed using 100 μL tips at 1150 V, 20 ms, and 2 pulse parameters. Cells were collected at 0 h, 4 h, 12 h, and 24 h postelectroporation, and the cleared lysate was transferred onto streptavidin Dynabeads (30 μl per sample; Invitrogen). After 3 washes with lysis buffer, the beads were boiled at 95 °C for 10 min and loaded onto the gel for WB.

### DCLRE1A protein production

The cleaved MBL-β-CASP domain of human SNM1A (aa 695–1040), both wild type and mutant forms (D745A, H746A), were cloned and inserted into the pcDNA3.1 plasmid vector with an added His_6_-tag on the C-terminus. The proteins were expressed in HEK293 cells for 48 h (15 cm dish x 4 for each protein, 20 µg per dish of plasmid DNA was transfected using jetPRIME transfection reagent (Polyplus)); then, the cells were washed twice with PBS and collected by centrifugation. Cells were lysed in 20 ml of lysis buffer (50 mM HEPES, pH 7.5, 500 mM NaCl, 20 mM imidazole, 5% glycerol, protease inhibitor cocktail (Roche), and 0.5 mM PMSF) using sonication (100% power, 5 sec on/5 sec off, Qsonica, Model CL-18) and centrifuged for 15 min at 15000 g and 4 °C. The supernatant was loaded on an FPLC column containing 1 ml of WorkBeads Ni-NTA resin (Bio-Works) equilibrated with the same lysis buffer. All chromatographic purification steps were performed using an Amersham Biosciences Akta Prime FPLC System. The column was subsequently washed with 10 ml of the same buffer, and the His-tagged protein was eluted using a gradient of 20 mM–1 M imidazole (10 ml). The eluted protein was concentrated using Amicon Ultra 10K Centrifugal Filters (Millipore) with exchange to storage buffer: 20 mM HEPES (pH 7.9), 50 mM KCl, 10 mM MgCl_2_, 0.5 mM DTT, 0.05% Triton X-100, and 20% glycerol. The purity of the obtained protein fractions was estimated using SDS‒PAGE with One-Step Blue^®^ Protein Gel Stain (Biotium) and by Western blotting with anti-His-tag antibodies.

### In vitro cleavage assays

SNM1A nuclease assays were performed in 10 µl of the following reaction buffer: 20 mM HEPES (pH 7.9), 50 mM KCl, 10 mM MgCl_2_, 0.5 mM DTT, 0.05% Triton X-100, 0.1 mg/ml BSA, and 5% glycerol with 0.5 µM fluorescently labeled DNA. The nuclease reaction was initiated by the addition of the protein, incubated at 37 °C for 2 h, and stopped by the addition of proteinase K (0.5 µg/µL final) and an incubation for another 15 min at 55 °C. The samples were mixed with loading buffer (80% formamide, 20% glycerol, and 10 mM EDTA) at a 1:0.5 ratio and loaded onto 20% denaturing PAG containing 7 M urea. Electrophoretically separated DNA fragments were detected at 526 nm using a Typhoon Imager (GE Healthcare).

### Coimmunoprecipitation of DCLRE1A with POLI

HEK293T_35CAG_ATN1 cells with stable Cas12a expression (2.5 × 10^6^) were transfected with 4 µg of a plasmid expressing FLAG-tagged DCLRE1A, mut DCLRE1A, and ΔUBZ-DCLRE1A (a gift from Peter J. McHugh and Roger A. Greenberg) and POLI (a gift from Justyna McIntyre) [48] using PEI. At 48 h posttransfection, 6x10^6^ cells were transfected with 800 pmol of g14 using Lipofectamine 2000 (Thermo Fisher Scientific), or bis(2-chloroethyl)amine hydrochloride (nitrogen mustard) (Thermo Fisher Scientific) was added immediately to serum-free medium at a final concentration of 1000 µM. At 12 h posttransfection or 12 h after nitrogen mustard addition, the cells were scraped in cold PBS supplemented with 1 mM PMSF. After centrifugation, the cells were lysed using 1.8 mL of nuclear lysis buffer (50 mM Tris-HCl (pH 7.5), 150 mM NaCl, 1% NP-40, 0.5% sodium deoxycholate, and 0.1% SDS) or lysis buffer for Benzonase treatment (50 mM Tris-HCl (pH 7.5), 150 mM NaCl, 1% NP-40, 0.5% sodium deoxycholate, 2mM MgCl2) and placed on ice for 30 min to ensure efficient lysis. To determine whether the interaction between DCLRE1A and POLI was DNA-dependent, 125 U of Benzonase Nuclease was added to the reaction mixture. Next, the lysates were centrifuged at 10,000 × g for 5 min to remove cellular debris and then incubated overnight with 50 μL of ANTI-FLAG M2 Magnetic Beads (Sigma‒Aldrich). After the overnight incubation, the beads were washed 3x with lysis buffer. Next, the beads were resuspended in 100 μL of lysis buffer and transferred to a clean tube. Finally, the beads were boiled at 95 °C for 10 min in elution buffer and then loaded onto the gel for WB.

#### Western blot

The samples were diluted in buffer containing 2-mercaptoethanol and denatured at 95 °C for 5 min. The samples were then loaded onto a 12% gel and separated at 135 V for approximately 2 h at 4 °C. The proteins were transferred to a nitrocellulose membrane (Sigma‒Aldrich) for 1 h at 100 V. The membranes were blocked with TBS containing 0.1% Tween-20% and 5% nonfat dry milk for 1 h at room temperature. Immunodetection was performed using antibodies against DCLRE1A, POLI, PCNA, ub-PCNA, and GAPDH (listed in Supplementary Table 5). The membranes were visualized using a chemiluminescent substrate for Western blotting, Westar Antares (Cyanagen).

### Rescue experiment

sgPOLI-1, sgPOLI-2, and sgDCLRE1A-1, and sgDCLRE1A-2 were cloned and inserted into the pSpCas9(BB)-2A-GFP (PX458) plasmid (#48138, Addgene) and used to knockout the POLI (RPEkoPOLI) and DCLRE1A (RPEkoDCLRE1A) genes, respectively. For double knockout (PREkoPOLIandDCLRE1A), RPEkoPOLI cells were treated with sgDCLRE1A-1 or sgDCLRE1A-2. Plasmids were delivered into cells via electroporation using the Neon transfection system. Briefly, ∼1 million cells were mixed with 2.5 μg of each plasmid in 100 μL tips, and electroporation parameters of 1350 V and 30 ms with one pulse were used. Next, one cell was sorted into each well of a 96-well plate based on the GFP signal. Cells were detached when they reached 50% confluence, and the knockout efficiency was verified using WB with anti-POLI and anti-DCLRE1A antibodies.

For the rescue experiments, cells were infected with a virus carrying 40 CAG repeats at an MOI of 0.3 and underwent puromycin selection until they reached a 100% BFP signal. RPEkoPOLI and RPEkoDCLRE1A cells were electroporated with 3 μg of each plasmid expressing Cas12a, POLI, or DCLRE1A. Plasmids expressing DCLRE1A, POLI, and Cas12a (3 µg of each plasmid) were delivered into RPEkoPOLIandDCLRE1A. Cells were sorted based on GFP and BFP signals, and on the next day, 100 pmol of g14 was delivered into the cells using Lipofectamine 2000 (Thermo Fisher Scientific). Cells were collected 48 h after g14 delivery, and DNA was isolated using a Genomic DNA Isolation Kit (Norgen, Biotek Corp.) according to the manufacturer’s instructions. Eighty nanograms of isolated DNA was used for nested PCR to amplify a DNA fragment containing the CAG repeats. Primers 171 and 172 were first used to amplify ca. 800 bp products. Next, the first PCR was 10x diluted for a second PCR with 1F and 2R primers using Phusion high-fidelity PCR master mix (Thermo Fisher Scientific). The purified PCR products were subjected to paired-end NGS analysis.

### Proteomics

#### RPE-Cas12a-BirA*_40CAG and RPE-dCas12a-BirA*_40CAG cell line generation

The BirA* sequence from the pcDNA3.1 mycBioID plasmid (#35700, Addgene) was cloned and inserted into the modified plasmid #188492 (see the Cloning and construct generation section). A D908A mutation was introduced into the Cas12a sequence to generate catalytically inactive dCas12a using the Gibson Assembly^®^ Cloning Kit with the primers Forward_fr1, Reverse_fr1, Forward_fr2, Reverse_fr2, Forward_fr3, and Reverse_fr3, as listed in Supplementary Table 2.

For lentivirus production, the plasmids with the expression of Cas12a-BirA* or dCas12a-BirA* were cotransfected with the packaging plasmids pCMV-VSV-G (#8454, Addgene) and pCMV-dR8.2dvpr (#8455, Addgene) into HEK 293T cells using the PEI transfection method. The medium containing lentivirus was collected on day 2 and passed through 0.45-μm filters.

RPE-1_40CAG cells (without BFP) were transduced with a lentivirus carrying either Cas12a-BirA* or dCas12a-BirA* at an MOI of 1 in the presence of polybrene (4 μg/mL). At 48 h postinfection, the cells were sorted into a 96-well plate (one cell per well) based on the BFP signal. After the cells reached 50% confluence, individual clones were tested for Cas12a/dCas12a expression and the effectiveness of biotinylation using an anti-HPRT antibody.

#### Protein extraction and immunoblotting

Total protein extracts were generated by lysing cells in PB buffer (BioShop). A total of 50 μg of protein was diluted in buffer containing 2-mercaptoethanol and incubated for 5 min at 95 °C. The samples were separated on a 10% Tris-acetate SDS-polyacrylamide gel and transferred to a nitrocellulose membrane (Sigma‒Aldrich). The membranes were blocked with TBS supplemented with 0.1% Tween-20% and 5% nonfat dry milk for 1 h at room temperature. The membranes were incubated with the primary monoclonal ANTI-FLAG^®^ M2 (Merck) antibody and secondary donkey anti-mouse IgG antibody (Jackson ImmunoResearch) in TBS supplemented with 0.1% Tween-20 and 5% nonfat dry milk. Immunoreactions were detected using WESTAR ANTARES (Cyanagen).

The cells were incubated with 50 μM biotin, and the procedure was performed as described above until membrane blocking to test the biotinylation activity. Afterward, the membranes were incubated overnight with HRP-streptavidin (Thermo Fisher Scientific).

#### BioID

RPE-Cas12a-BirA*_40CAG and RPE-dCas12a-BirA*_40CAG cells (0.7 × 10^6^) were seeded in 10 cm^2^ dishes in DMEM-F12 supplemented with 10% FBS and 50 μM biotin. Twenty-four hours after the biotin treatment, the cells were lipofected with 16 μM sg14 and collected 12 h after the editing process was initiated. For this analysis, 4 biological replicates were performed. Next, the cells were washed 7x with cold PBS, collected by scraping in cold PBS supplemented with 1 mM PMSF and centrifuged for 5 min at 800 × *g* at 4 °C. The pelleted cells were incubated in 0.5 ml of cellular lysis buffer (5 mM PIPES (pH 8), 85 mM KCl, and 0.1% Triton-1000x) on ice for 5 min and centrifuged for 5 min at 800 × *g* at 4 °C. Next, the cells were lysed with 0.6 ml of nuclear lysis buffer (50 mM Tris buffer (pH 7.5), 0.5% sodium deoxycholate, 150 mM NaCl, 1% NP40, and 0.1% SDS) and incubated on ice for 15 min. The cellular lysates were sonicated using a BioRuptor Pico sonicator (Diagenode; 2 cycles, 30 s ON, 30 s OFF), and the supernatants were collected and transferred to new tubes. The lysates were centrifuged for 5 min at 12500 × *g* at 4 °C to remove cellular debris, transferred to new tubes (LoProtein, Eppendorf), and stored at -80 °C.

Two hundred microliters of neutravidin agarose beads (Thermo Fisher Scientific) suspended in a storage solution were centrifuged for 1 min at 500 × *g*, after which the supernatant was removed. The beads were washed 3x with nuclear lysis buffer, and the lysates were incubated with the beads for 90 min at 4 °C with gentle rotation. Next, the beads were centrifuged for 1 min at 500 × *g*, washed 3x with PBS and 5x with HEPES (pH 8.0), suspended in 100 μl of HEPES (pH 8.0) and stored at 4 °C until the mass spectrometry analysis was performed.

### Mass spectrometry

#### Liquid chromatography‒mass spectrometry measurement

Before the liquid chromatography‒mass spectrometry (LC‒MS/MS) measurements were performed, the peptide fractions were resuspended in 0.1% trifluoroacetic acid (TFA) and 2% acetonitrile in water. Chromatographic separation was performed on an Easy-Spray Acclaim PepMap column (50 cm length × 75 µm inner diameter; Thermo Fisher Scientific) at 55 °C by applying 75 min of gradient acetonitrile concentrations in 0.1% aqueous formic acid at a flow rate of 300 nl/min. An UltiMate 3000 nano-LC system was coupled to a Q Exactive HF-X mass spectrometer via an easy-spray source (all Thermo Fisher Scientific). The Q Exactive HF-X was operated in TMT mode with survey scans acquired at a resolution of 60,000 at m/z 200. Up to 15 of the most abundant isotope patterns with charges of 2–5 from the survey scan were selected with an isolation window of 0.7 m/z and fragmented by higher-energy collision dissociation with normalized collision energies of 32, while the dynamic exclusion was set to 35 s. The maximum ion injection times for the survey scan and dual MS (MS/MS) scans (acquired with a resolution of 45,000 at m/z 200) were 50 and 96 ms, respectively. The ion target value for MS was set to 3 × 10^6, that for MS/MS was set to 1 × 10^5, and the minimum AGC target was set to 1 × 10^3.

#### Data Processing

The data were processed with MaxQuant v. 1.6.17.0, and the peptides identified in the MS/MS spectra were searched against the UniProt Human Reference Proteome (UP000005640) using the built-in Andromeda search engine. Reporter ion MS2-based quantification was applied with a reporter mass tolerance = 0.003 Da and a minimum reporter PIF = 0.75. Cysteine carbamidomethylation was set as a fixed modification, and methionine oxidation, glutamine/asparagine deamination, and protein N-terminal acetylation were set as variable modifications. For in silico digests of the reference proteome, cleavage of arginine or lysine followed by any amino acid was allowed (trypsin/P), and up to two missed cleavages were allowed. The FDR was set to 0.01 for the peptides, proteins and sites. Match between runs was enabled. Other parameters were used as preset in the software. Reporter intensity-corrected values for protein groups were loaded into Perseus v. 1.6.10. Standard filtering steps were applied to clean the dataset: reverse (matched to a decoy database), identified only by site, and potential contaminants (from a list of commonly occurring contaminants included in MaxQuant) were removed. Reporter intensity-corrected values were log2 transformed, and protein groups with values across all samples were retained. The values were normalized by median subtraction within the TMT channels. Student’s t test (2-sided, permutation-based FDR = 0.01, S0 = 0.1, n = 4) was performed to determine the proteins that were differentially regulated. The obtained data were deposited in the ProteomeXchange Consortium via the PRIDE partner repository with the dataset identifiers PXD070360.

### Cloning and construct generation

A modified AsCas12a plasmid (#188492, Addgene) was used to generate constructs with repair factors. First, using KpnI and NheI, the fragment of the plasmid containing the Cas9 scaffold was removed. Next, the BFP sequence was cloned at newly generated EcoRI restriction sites following P2A cloning at the BamHI and EcoRI restriction sites. Additionally, the linker (L) was cloned at the AfeI restriction site between Cas12a and the gene sequence of interest. The genes of interest were subsequently cloned using the Gibson Assembly^®^ Cloning Kit. Phusion Flash High-Fidelity PCR Master Mix and the primers pl_F and pl_R were used to amplify the entire plasmid sequence. The primers used for Gibson cloning, which amplified DCLRE1A, FANCG, SAP, and MLR, are listed in Table 1. A plasmid expressing Cas12a-PALB2 was synthesized in GeneScript.

### DNA amplification and targeted deep sequencing

Genomic DNA was extracted using a Genomic DNA Isolation Kit (Norgen, Biotek Corp.) according to the manufacturer’s instructions and quantified using a spectrophotometer/fluorometer (DeNovix). The DNA obtained from cell cultures transfected with the RNP complexes was amplified by nested PCR using Phusion Flash High-Fidelity PCR Master Mix (Thermo Fisher Scientific). The first amplification of DNA from HEK293T_35CAG_ATN1 cells and hybrids was performed with the primers ATN1_F and ATN1_R as follows: initial denaturation at 98 °C for 1 min; 25 cycles at 98 °C for 5 s, 63.2 °C for 5 s, and 72 °C for 15 s; and a final elongation at 72 °C for 1 min. PCR products were diluted 10 × and used as a template for the second amplification performed with the primers End_1F and End_1R as follows: initial denaturation at 98 °C for 1 min; 30 cycles at 98 °C for 5 s, 65 °C for 5 s, and 72 °C for 15 s; and a final elongation at 72 °C. Nested PCR was performed using Phusion Flash High-Fidelity PCR Master Mix (Thermo Fisher Scientific) to analyze the pure contraction/aberrant contraction ratio and detect inverted repeats. The first amplification was performed with primers 171 and 172 as follows: initial denaturation at 98 °C for 1 min; 25 cycles at 98 °C for 5 s, 68 °C for 5 s, and 72 °C for 20 s; and a final elongation at 72 °C for 1 min. PCR products were diluted 10 × and used as a template for the second amplification performed with primers 1F and 2R as follows: initial denaturation at 98 °C for 1 min; 25 cycles at 98 °C for 5 s, 63.1 °C for 5 s, and 72 °C for 15 s; and a final elongation at 72 °C. The sequences of the specific primers are listed in Additional file 2, Table S1. The PCR products were used for library generation and next-generation sequencing, which were performed by Novogene (UK). The sequencing libraries were generated using the NEBNext Ultra II DNA Library Prep Kit for Illumina (New England Biolabs) according to the manufacturer’s protocol. Libraries were sequenced using the Illumina NovaSeq 6000 platform with the following settings: 250 bp paired-end reads and 1 million reads per sample. The obtained data were deposited in the Sequence Read Archive (SRA, accession number: PRJNA1347391).

### Bioinformatic analysis

The analysis was performed by ideas4biology Ltd. Briefly, the overall data quality was tested with FastQC version 0.11.5. Adapters were removed using a standard adapter sequence set from the bbduk2 package. Reads with a quality less than 5 were discarded. The reads were merged using the bbmerge script from the BBMap package. After merging, the reads were mapped against the *ATN1* gene sequence using the BBMap global aligner. Mapping quality was assessed using qualimap v.2.2.2-dev. Aligned reads were filtered using the -F260 and -q 30 flags. The sequences were clustered using the cd-hit application [49]. Sequences with the same length and at least 95% sequence similarity were merged into clusters. The bam files containing unique sets of reads were converted to pairwise alignments using the sam2pairwise program. We used Tandem Repeats Finder v409 to locate and display tandem repeats in DNA sequences. The recommended parameters were used except for the minimum alignment score, which was set to 10 to allow for shorter hits. The results were filtered for repeats containing only combinations of the letters C, A, and G. Reads for each “CAG” repeat length were counted, and the ratio of the number of reads containing a given “CAG” length to all reads was calculated.

### Generation of the HEK-Flp-In_T-Rex_64CAG cell line

Fragments of the *ATN1* gene with 64 CAG repeats were cloned and inserted into the pcDNA5/FRT/TO vector (Invitrogen). This vector was integrated into the genome via Flp recombinase-mediated DNA recombination at the FRT site. The sequences of the DNA oligonucleotides used for cloning are provided in Table 7. In Supplementary Material 2, the fragments of the integrated sequence are listed. The pcDNA5/FRT/TO-based expression construct and pOG44 vector were cotransfected at a 1:9 ratio into Flp-In T-REx-293 host cells using Lipofectamine 2000 according to the manufacturer’s protocol. Hygromycin-resistant monoclones containing stably integrated expression cassettes were selected using 100 μg/ml hygromycin B according to the manufacturer’s guidelines. The medium was replaced every 3 to 4 days to eliminate dead cells. After 2 to 3 weeks, individual hygromycin-resistant colonies were clonally selected through manual selection under an EVOS microscope. The individual colonies were further expanded and stored in liquid nitrogen using the same medium supplemented with 10% DMSO. The expression of recombinant proteins was screened following cell induction with doxycycline (Sigma‒Aldrich) using immunoblotting.

### Measurement of the in-frame/out-of-frame ratio in the HEK-Flp-In_T-Rex_64CAG cells

Twenty-four hours before transfection, 4x10^5^ HEK-Flp-In_T-Rex_64CAG cells were seeded into one well of a 6-well plate. Then, 2 µg of plasmid expressing Cas12a-repair factors were introduced into cells using Lipofectamine 2000. At 48 h after plasmid transfection, 1x10^5^ HEK-Flp-In_T-Rex_64CAG cells were seeded onto a 12-well plate and transfected with 100 pmol of g14. At 48 h after g14 delivery, the cells were treated with 1 µg/ml doxycycline (Sigma‒Aldrich), and the cells were subjected to a cytometric analysis 24 h after doxycycline induction. The cytometric analysis was performed on a Guava easyCyte^TM^ 12HT. Cells were characterized by two nonfluorescent parameters, forward scatter (FSC) and side scatter (SSC), and two fluorescence parameters: green fluorescence from GFP using a 488 nm laser (Ex/Em 488/507 nm) and blue fluorescence from BFP using a 405 nm laser (Ex/Em 381/445 nm). The data were analyzed with GuavaSoft™ 4.0. Pure contractions were estimated by calculating the number of cells in the GFP/BFP channel (pure contractions) compared to events in the BFP channel alone (aberrant contractions and nonedited cells).

### Statistical analysis

Statistical analysis was performed using GraphPad Prism v. 5.0 software (GraphPad). The data were analyzed using an unpaired t test (∗p < 0.03, ∗∗p < 0.002, ∗∗∗p < 0.0002, ∗∗∗∗p < 0.0001), with an arbitrary value of 1 assigned to the unmodified RPE1 cells (Supplementary Figure 7 and Supplementary Figure 16) or to the cells treated with non-targeting gRNAs (Figures 4D, 5B) or knockout of specific gene (Figure 4E, Supplementary Figure 14) or BIRA (Figure 6D). Statistical analysis of the data in Figure 3A was performed using the Mann‒Whitney and Kolmogorov‒Smirnov tests. Asterisks denote the levels of statistical significance: **** p ≤ 0001.

## Acknowledgments

We thank Grzegorz Figura for his support during the cloning process; Konrad Kuczynski for his help during the microscopic analysis; Marta Kazimierska for her support in capillary gel electrophoresis and microscopic analysis; Remigiusz Serwa from the Proteomics Core Facility IMol, Warsaw, for performing mass spectrometry; Peter J. McHugh and Lonnie P. Swift from the Department of Oncology, MRC-Weatherall Institute of Molecular Medicine, University of Oxford; and Tianpeng Zhang and Roger A. Greenberg from the Department of Cancer Biology, Penn Center for Genome Integrity, Basser Center for BRCA, Perelman School of Medicine, University of Pennsylvania, for kindly providing the plasmid expressing DCLRE1A. We thank Justyna McIntyre from the Laboratory of Mutagenesis and DNA Damage Tolerance, Institute of Biochemistry and Biophysics, Polish Academy of Sciences, for providing the plasmid expressing POLI. We thank Magdalena Trybus from the Laboratory of Single-Cell Analyses IBCH PAS for performing the flow cytometry experiments and cell sorting. The equipment used was sponsored in part by the Centre for Preclinical Research and Technology (CePT), a project cosponsored by the European Regional Development Fund and the Innovative Economy, as well as the National Cohesion Strategy of Poland. The flow cytometry data collected in this study were acquired using the infrastructure developed under the project NEBI—National Imaging Centre for Biological and Biomedical Sciences, POIR.04.02.00-00-C004/19, co-financed through the European Regional Development Fund (ERDF) in the frame of the Smart Growth Operational Programme 2014-2020 (Measure 4.2 Development of modern research infrastructure of the science sector). A L’Oréal-UNESCO For Women in Science fellowship supported M.D. throughout this project.

## Data availability

All data generated or analyzed during this study are included in this published article, its supplementary information files, and publicly available repositories. The targeted sequencing data were deposited in the Sequence Read Archive under the accession number: PRJNA1347391. The mass spectrometry proteomics data were deposited in the ProteomeXchange Consortium via the PRIDE partner repository with the dataset identifiers PXD070360.

## References

[1] C. Depienne and J. L. Mandel, “30 years of repeat expansion disorders: What have we learned and what are the remaining challenges?,” Am. J. Hum. Genet., vol. 108, no. 5, pp. 764–785, May 2021.

[2] E. L. Bunting, J. Hamilton, and S. J. Tabrizi, “Polyglutamine diseases,” Curr. Opin. Neurobiol., vol. 72, pp. 39–47, Feb. 2022.

[3] C. Pressl et al., “Selective vulnerability of layer 5a corticostriatal neurons in Huntington’s disease,” Neuron, vol. 112, no. 6, pp. 924–941.e10, Mar. 2024.

[4] K. Mätlik et al., “Cell-type-specific CAG repeat expansions and toxicity of mutant Huntingtin in human striatum and cerebellum,” Nat. Genet., vol. 56, no. 3, pp. 383–394, Mar. 2024.

[5] N. Wang et al., “Distinct mismatch-repair complex genes set neuronal CAG-repeat expansion rate to drive selective pathogenesis in HD mice,” Cell, vol. 188, no. 6, Feb. 2025.

[6] J. M. Lee et al., “CAG Repeat Not Polyglutamine Length Determines Timing of Huntington’s Disease Onset,” Cell, vol. 178, no. 4, pp. 887–900.e14, Aug. 2019.

[7] G. F. Richard, D. Viterbo, V. Khanna, V. Mosbach, L. Castelain, and B. Dujon, “Highly specific contractions of a single CAG/CTG trinucleotide repeat by TALEN in yeast,” PLoS One, vol. 9, no. 4, Apr. 2014.

[8] V. Mosbach, L. Poggi, D. Viterbo, M. Charpentier, and G. F. Richard, “TALEN-Induced Double-Strand Break Repair of CTG Trinucleotide Repeats,” Cell Rep., vol. 22, no. 8, pp. 2146–2159, Feb. 2018.

[9] F. K. Ekman, D. S. Ojala, M. M. Adil, P. A. Lopez, D. V. Schaffer, and T. Gaj, “CRISPR-Cas9-Mediated Genome Editing Increases Lifespan and Improves Motor Deficits in a Huntington’s Disease Mouse Model,” Mol. Ther. Nucleic Acids, vol. 17, p. 829, Sep. 2019.

[10] P. Sledzinski, M. Nowaczyk, M. I. Smielowska, and M. Olejniczak, “CRISPR/Cas9-induced double-strand breaks in the huntingtin locus lead to CAG repeat contraction through DNA end resection and homology-mediated repair,” BMC Biol., vol. 22, no. 1, p. 282, Dec. 2024.

[11] C. Cinesi, L. Aeschbach, B. Yang, and V. Dion, “Contracting CAG/CTG repeats using the CRISPR-Cas9 nickase,” Nat. Commun. *2016* 71, vol. 7, no. 1, pp. 1–10, Nov. 2016.

[12] D. Mittelman et al., “Zinc-finger directed double-strand breaks within CAG repeat tracts promote repeat instability in human cells,” vol. 106, no. 24, pp. 9607–9612, 2009.

[13] C. S. Casas-Delucchi, M. Daza-Martin, S. L. Williams, and G. Coster, “The mechanism of replication stalling and recovery within repetitive DNA,” Nat. Commun., vol. 13, no. 1, Dec. 2022.

[14] E. J. Polleys, I. Del Priore, J. E. Haber, and C. H. Freudenreich, “Structure-forming CAG/CTG repeats interfere with gap repair to cause repeat expansions and chromosome breaks,” Nat. Commun., vol. 14, no. 1, Dec. 2023.

[15] R. Ceccaldi, B. Rondinelli, and A. D. D’Andrea, “Repair Pathway Choices and Consequences at the Double-Strand Break,” Trends Cell Biol., vol. 26, no. 1, p. 52, Jan. 2015.

[16] B. Zetsche et al., “Cpf1 is a single RNA-guided endonuclease of a class 2 CRISPR-Cas system,” Cell, vol. 163, no. 3, pp. 759–771, Oct. 2015.

[17] D. C. Swarts and M. Jinek, “Cas9 versus Cas12a/Cpf1: Structure-function comparisons and implications for genome editing,” Wiley Interdiscip. Rev. RNA, vol. 9, no. 5, Sep. 2018.

[18] I. Strohkendl, F. A. Saifuddin, J. R. Rybarski, I. J. Finkelstein, and R. Russell, “Kinetic Basis for DNA Target Specificity of CRISPR-Cas12a,” Mol. Cell, vol. 71, no. 5, pp. 816–824.e3, Sep. 2018.

[19] B. P. Kleinstiver et al., “Engineered CRISPR-Cas12a variants with increased activities and improved targeting ranges for gene, epigenetic and base editing,” Nat. Biotechnol., vol. 37, no. 3, pp. 276–282, Mar. 2019.

[20] Z. Gao, M. Fan, A. T. Das, E. Herrera-Carrillo, and B. Berkhout, “Extinction of all infectious HIV in cell culture by the CRISPR-Cas12a system with only a single crRNA,” Nucleic Acids Res., vol. 48, no. 10, pp. 5527–5539, 2020.

[21] H. Zhang et al., “An engineered xCas12i with high activity, high specificity, and broad PAM range,” Protein Cell, vol. 14, no. 7, p. 540, Jul. 2022.

[22] L. A. Gilbert et al., “CRISPR-mediated modular RNA-guided regulation of transcription in eukaryotes,” Cell, vol. 154, no. 2, p. 442, Jul. 2013.

23. “Introduction - Handbook of Biological Statistics.” [Online]. Available: https://www.biostathandbook.com/. [Accessed: 17-Oct-2025].

[24] T. Iyama et al., “CSB interacts with SNM1A and promotes DNA interstrand crosslink processing,” Nucleic Acids Res., vol. 43, no. 1, p. 247, Sep. 2014.

[25] J. Michl, J. Zimmer, and M. Tarsounas, “Interplay between Fanconi anemia and homologous recombination pathways in genome integrity,” EMBO J., vol. 35, no. 9, pp. 909–923, May 2016.

[26] S. Hashimoto, H. Anai, and K. Hanada, “Mechanisms of interstrand DNA crosslink repair and human disorders,” Genes Environ., vol. 38, no. 1, p. 9, 2016.

[27] T. Zhang, Y. Rawal, H. Jiang, Y. Kwon, P. Sung, and R. A. Greenberg, “Break induced replication orchestrates resection dependent template switch,” Nature, vol. 619, no. 7968, p. 201, Jul. 2023.

[28] B. Buzon, R. Grainger, S. Huang, C. Rzadki, and M. S. Junop, “Structure-specific endonuclease activity of SNM1A enables processing of a DNA interstrand crosslink,” Nucleic Acids Res., vol. 46, no. 17, p. 9057, Sep. 2018.

[29] L. P. Swift et al., “SNM1A is crucial for efficient repair of complex DNA breaks in human cells,” Nat. Commun., vol. 15, no. 1, p. 5392, Dec. 2024.

[30] A. T. Wang et al., “Human SNM1A and XPF-ERCC1 collaborate to initiate DNA interstrand cross-link repair,” Genes Dev., vol. 25, no. 17, pp. 1859–1870, Sep. 2011.

[31] H. T. Baddock, Y. Yosaatmadja, J. A. Newman, C. J. Schofield, O. Gileadi, and P. J. McHugh, “The SNM1A DNA repair nuclease,” DNA Repair (Amst)., vol. 95, p. 102941, Nov. 2020.

[32] A. Tissier, J. P. McDonald, E. G. Frank, and R. Woodgate, “polι, a remarkably error-prone human DNA polymerase,” Genes Dev., vol. 14, no. 13, p. 1642, Jul. 2000.

[33] C. J. Sakofsky, S. Ayyar, A. K. Deem, W. H. Chung, G. Ira, and A. Malkova, “Translesion polymerases drive microhomology-mediated break-induced replication leading to complex chromosomal rearrangements,” Mol. Cell, vol. 60, no. 6, p. 860, Dec. 2015.

[34] B. J. Payliss, A. Patel, A. C. Sheppard, and H. D. M. Wyatt, “Exploring the Structures and Functions of Macromolecular SLX4-Nuclease Complexes in Genome Stability,” Front. Genet., vol. 12, p. 784167, Nov. 2021.

[35] X. Xu et al., “Structure specific DNA recognition by the SLX1–SLX4 endonuclease complex,” Nucleic Acids Res., vol. 49, no. 13, p. 7740, Jul. 2021.

[36] H. Baechtold, M. Kuroda, J. Sok, D. Ron, B. S. Lopez, and A. T. Akhmedov, “Human 75-kDa DNA-pairing Protein Is Identical to the Pro-oncoprotein TLS/FUS and Is Able to Promote D-loop Formation,” J. Biol. Chem., vol. 274, no. 48, pp. 34337–34342, Nov. 1999.

[37] A. M. Couturier et al., “Roles for APRIN (PDS5B) in homologous recombination and in ovarian cancer prediction,” Nucleic Acids Res., vol. 44, no. 22, p. 10879, Dec. 2016.

[38] M. L. Nicolette et al., “Mre11–Rad50–Xrs2 and Sae2 promote 5’ strand resection of DNA double-strand breaks,” Nat. Struct. Mol. Biol., vol. 17, no. 12, p. 1478, Dec. 2010.

[39] L. Poggi, L. Emmenegger, S. Descorps-Declère, B. Dumas, and G. F. Richard, “Differential efficacies of Cas nucleases on microsatellites involved in human disorders and associated off-target mutations,” Nucleic Acids Res., vol. 49, no. 14, p. 8120, Aug. 2021.

[40] J. A. Hussmann et al., “Mapping the Genetic Landscape of DNA Double-strand Break Repair,” Cell, vol. 184, no. 22, p. 5653, Oct. 2021.

[41] T. Sijacki et al., “The DNA-damage kinase ATR activates the FANCD2-FANCI clamp by priming it for ubiquitination,” Nat. Struct. Mol. Biol., vol. 29, no. 9, p. 881, Sep. 2022.

[42] P. Alcón et al., “FANCD2–FANCI surveys DNA and recognizes double- to single-stranded junctions,” Nature, vol. 632, no. 8027, p. 1165, Aug. 2024.

[43] P. Tonzi and T. T. Huang, “Role of Y-family translesion DNA polymerases in replication stress: Implications for new cancer therapeutic targets,” DNA Repair, vol. 78. Elsevier B.V., pp. 20–26, 01-Jun-2019.

[44] S. K. Martin and R. D. Wood, “DNA polymerase ζ in DNA replication and repair,” Nucleic Acids Res., vol. 47, no. 16, p. 8348, Mar. 2019.

[45] D. Freedman and P. Diaconis, “On the histogram as a density estimator:L2 theory,” Zeitschrift für Wahrscheinlichkeitstheorie und Verwandte Gebiete, vol. 57, no. 4, pp. 453–476, 1981.

[46] W. Li et al., “MAGeCK enables robust identification of essential genes from genome-scale CRISPR/Cas9 knockout screens,” Genome Biol., vol. 15, no. 12, p. 554, Dec. 2014.

[47] T. Hart et al., “High-Resolution CRISPR Screens Reveal Fitness Genes and Genotype-Specific Cancer Liabilities,” Cell, vol. 163, no. 6, pp. 1515–1526, Dec. 2015.

[48] J. McIntyre et al., “DNA polymerase ι is acetylated in response to SN2 alkylating agents,” Sci. Rep., vol. 9, no. 1, p. 4789, Dec. 2019.

[49] W. Li and A. Godzik, “Cd-hit: a fast program for clustering and comparing large sets of protein or nucleotide sequences,” Bioinformatics, vol. 22, no. 13, pp. 1658–1659, Jul. 2006.

