## Supplemental Figures for "DCLRE1A orchestrates CAG repeat contraction following Cas12a-induced DNA breaks"

### Supplementary Figures

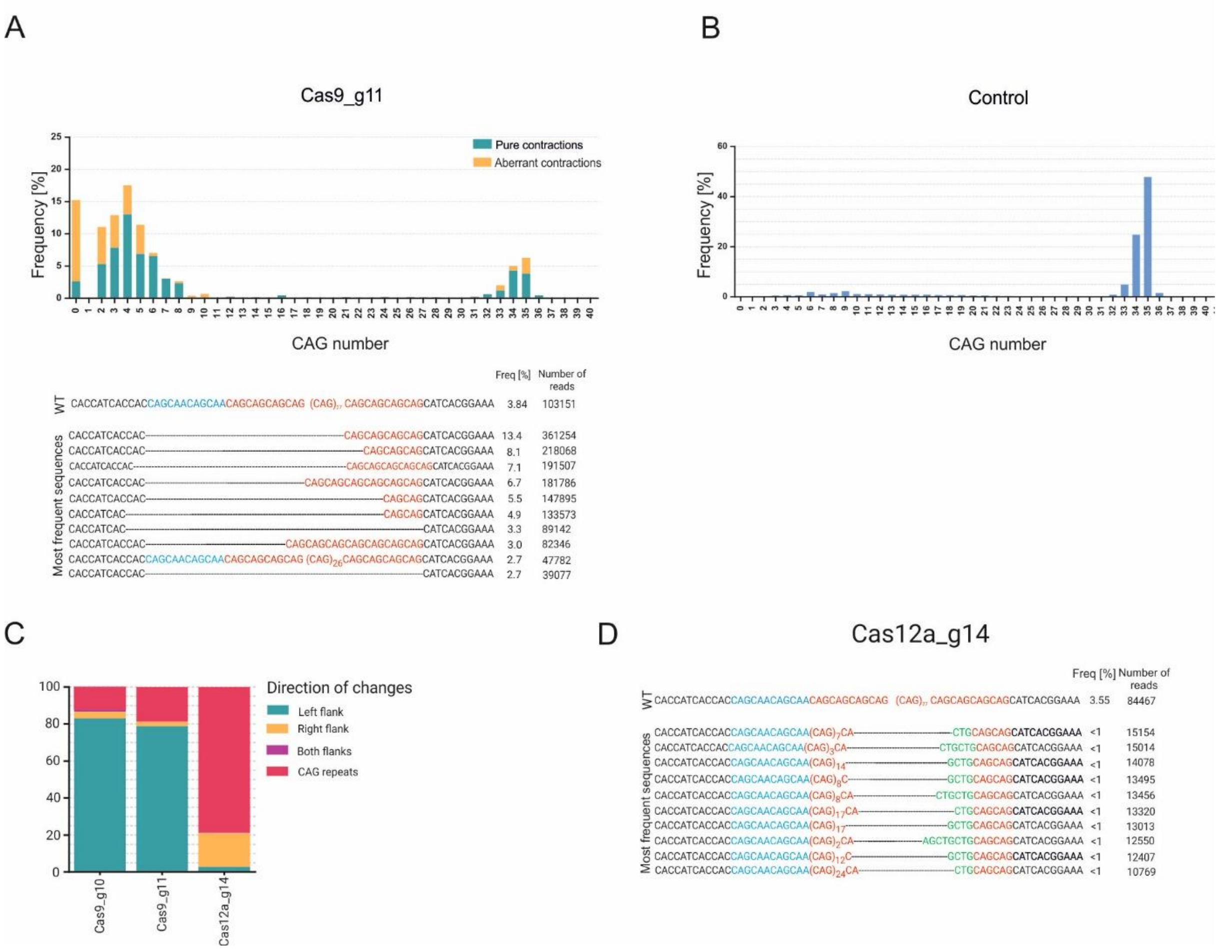


**Figure S1.** Repair outcomes after Cas9- and Cas12a-induced DSBs. **A.** Deep sequencing data revealed a long deletion of the CAG repeats after the use of Cas9_g11. The top ten sequences from the NGS are included in the graph below. The CAG repeats and CAG/CAA triplets located 5' upstream of pure CAG repeats are highlighted in red and blue, respectively. Deletions of the CAG repeats after Cas9 cleavage are shown as dashed lines. **B.** NGS results for non-edited samples (Control). **C.** The direction of changes following Cas9- and Cas12a-induced cleavage within the CAG repeats. **D.** The top ten sequences from the NGS data included inverted repeats and CTG mutant variants (highlighted in green).

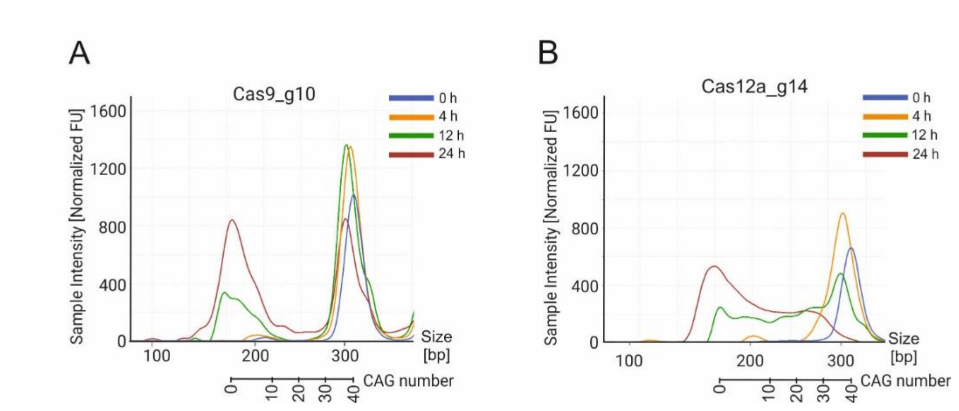


**Figure S2.** Dynamics of the contraction process. Capillary gel electrophoresis revealed differences in the dynamics of CAG repeat contraction following the delivery of Cas9_g10 (**A**) and Cas12a_g14 (**B**) into the RPE-1_40CAG cell line. Cells were collected at 0, 4, 12, and 24 h after the initiation of the editing process.

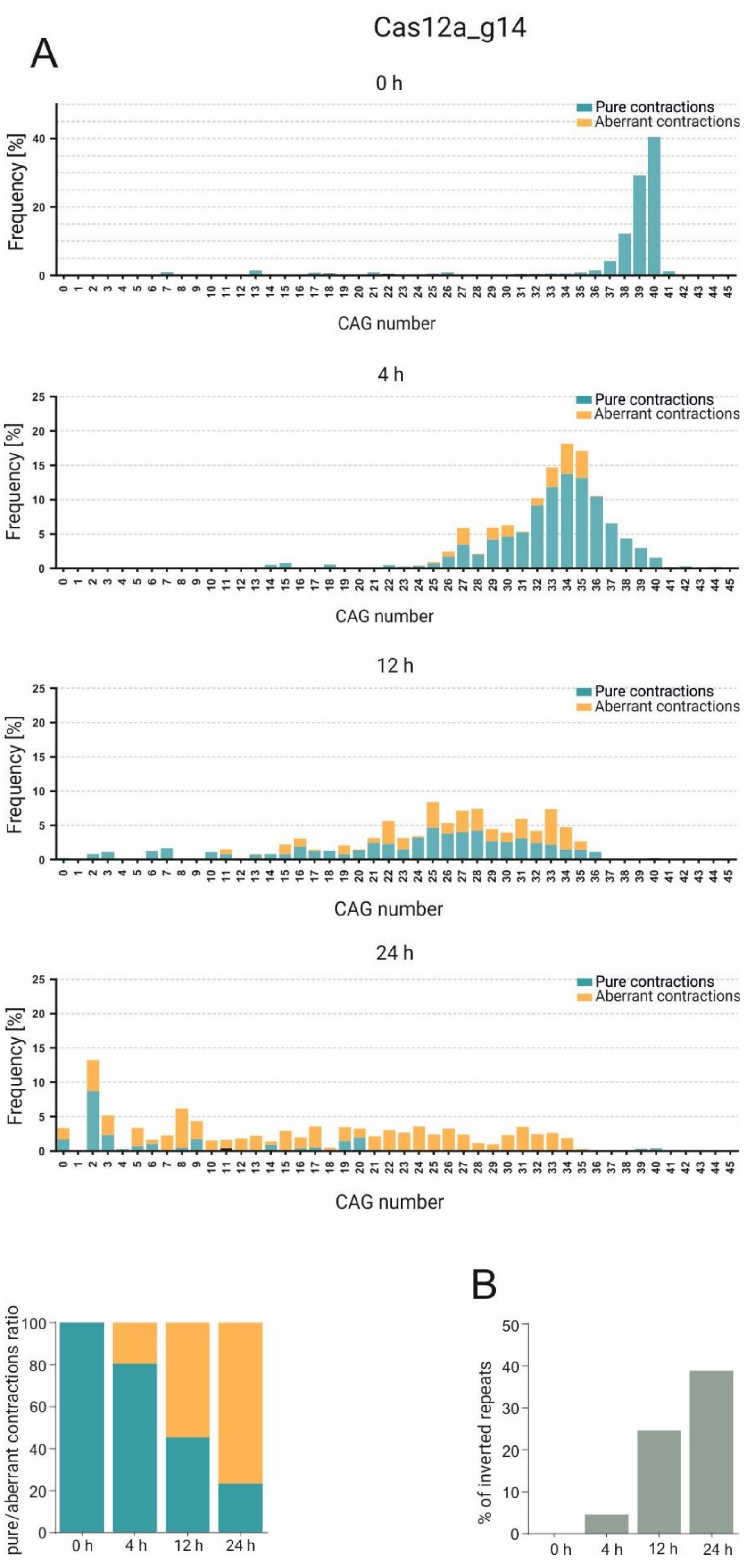


**Figure S3.** The dynamics of the CAG contraction process following Cas12a-induced cleavage. **A**. Deep sequencing data from RPE-1_40CAG cells treated with Cas12a_g14 revealed differences in the pure/aberrant contraction ratio and contraction pattern during editing. Cells were collected at 0, 4, 12, and 24 h after Cas12a_g14 delivery. **B**. Frequencies of inverted repeats at different time points.

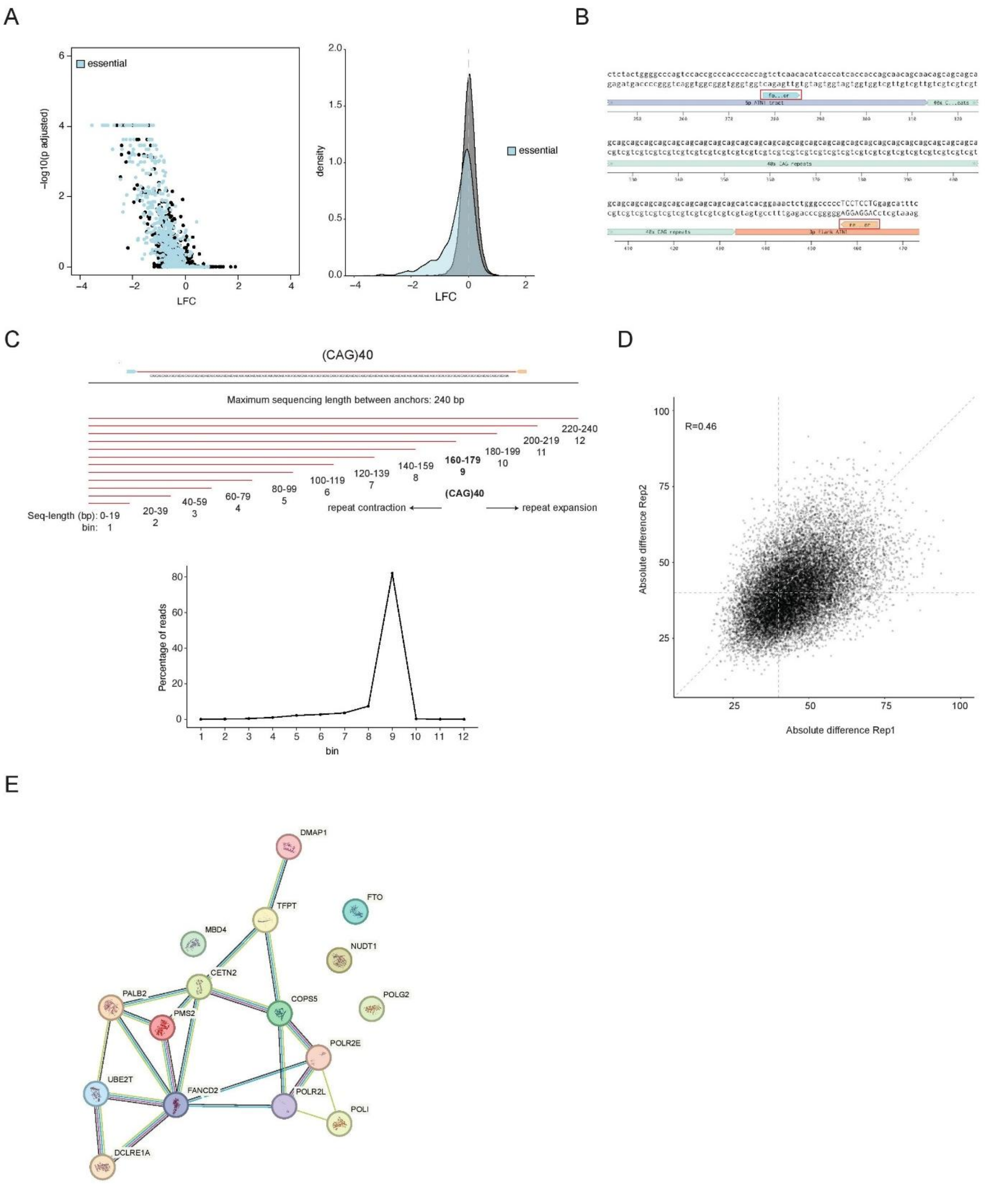


**Figure S4.** Quality control and results of the CRISPRi screen. **A.** Scatterplot showing the negative LFCs and the negative false discovery rate (fdr) comparing the plasmid library with the endpoint sample of the screen (Rep1), which was not treated with Cas12a (left panel). Density plot showing the log2(fold changes) (LFCs) of all genes, comparing the plasmid library with one endpoint sample of the screen (Rep1). Common essential genes are displayed in blue and show a lower LFC and lower negative fdr, demonstrating the successful knockdown of genes in the screen. **B.** Schematic of the (CAG)40 constructs in the library with the fwd (blue) and reverse (orange) anchor sequences used to determine sequence length between them marked. **C.** Schematic of the binning of sequences of different lengths into 12 bins. Bin 12–10 contain reads resulting from expansion events, whereas bin 8 and smaller contain sequences resulting from repeat contraction events. After all the reads were binned, the percentage of reads in each bin was calculated. **D.** Correlation plot of the absolute difference in each gene in comparison to the nontargeting distribution in replicate 1 and replicate 2 of the screens. Pearson's correlation coefficients between the replicates are shown (R = 0.46). **E.** STRING analysis of all 16 DNA repair genes identified to significantly change the CAG repeat pattern after Cas12a editing.

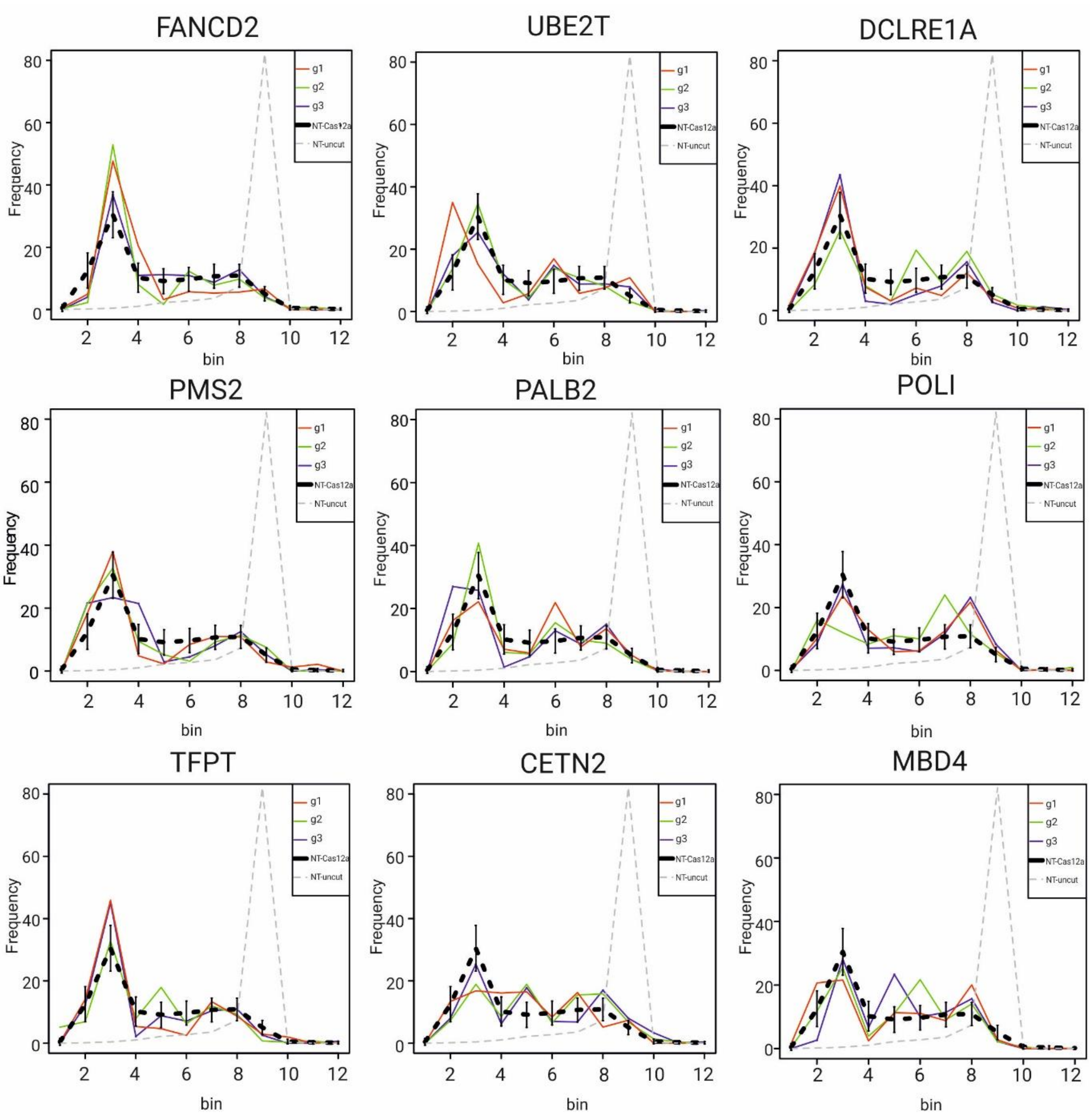


**Figure S5.** Frequency plots of DNA repair-related genes extracted from the CRISPRi screen. Sequence length distribution is shown for three gRNAs (g1, g2, and g3). NT-Cas12a indicates cells infected with a virus carrying a nontargeting gRNA (NT) and experiencing Cas12a-mediated DSBs. NT-uncut indicates cells infected with a virus carrying a nontargeting gRNA that remains uncut.

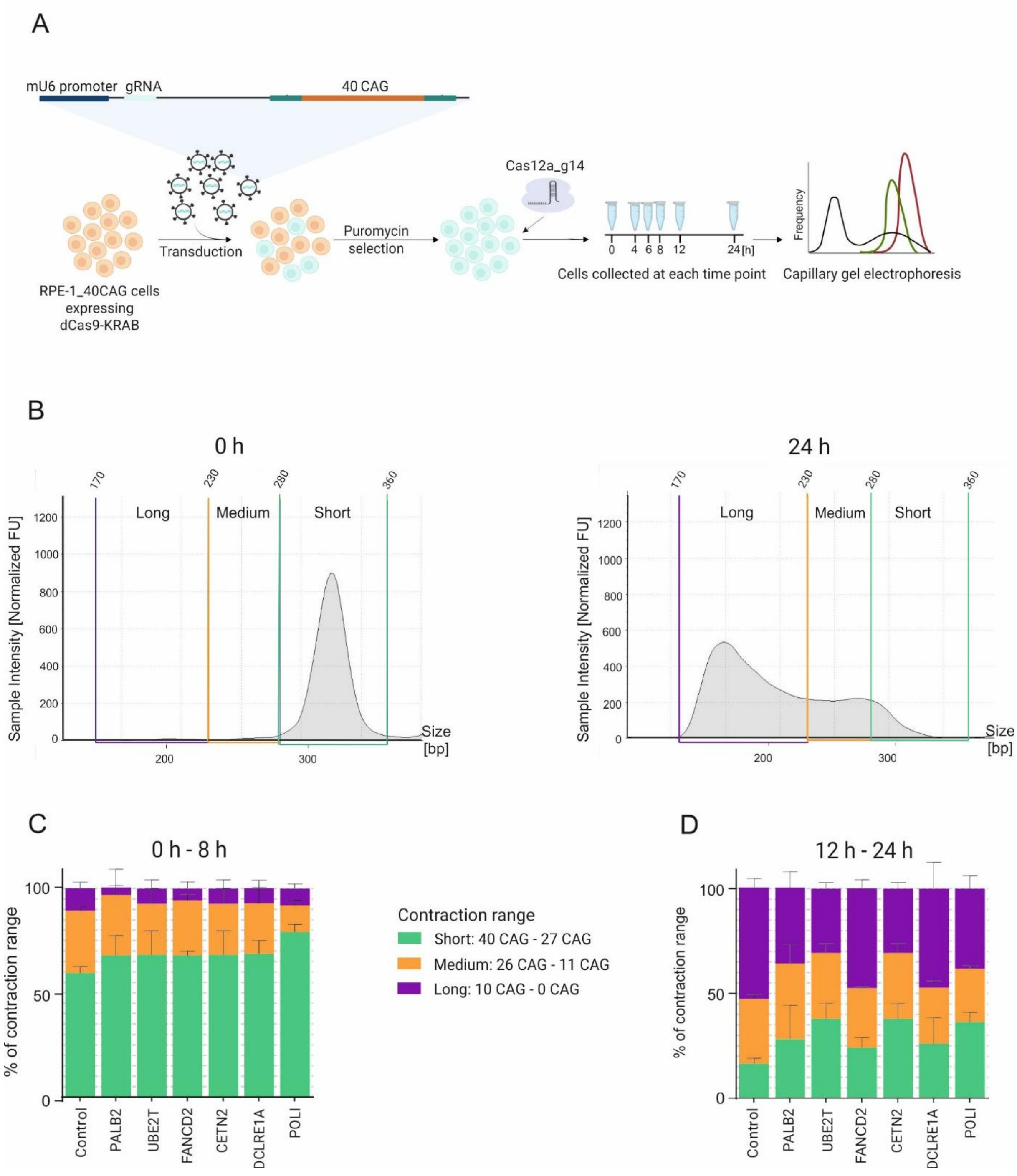


**Figure S6.** Dynamics of the contraction process. **A.** Schematic representation of the analysis of the contraction pattern in RPE-1_40CAG cells following the knockdown of candidate proteins chosen from the CRISPRi screen. RPE-1 cells stably expressing dCas9-KRAB were infected with viruses carrying gRNAs targeting the TSS of PALB2, UBE2T, FANCD2, CETN2, DCLRE1A, and POLI, as well as the nontargeting gRNA. Cells were treated with Cas12a_g14 and collected at different time points (0 h, 4 h, 6 h, 8 h, 12 h, and 24 h). Capillary gel electrophoresis was used to determine the contraction patterns. **B.** The amplified fragment containing 40 CAG repeats was divided into short (40–27 CAG repeats), medium (26–11 CAG repeats), and long (10–0 CAG repeats) contraction ranges. **C-D.** The concentrations of shortened variants at different time points during the contraction process were calculated for each range; next, the average concentrations for the short, medium, and long ranges at 0 h–8 h and 12 h–24 h were calculated.

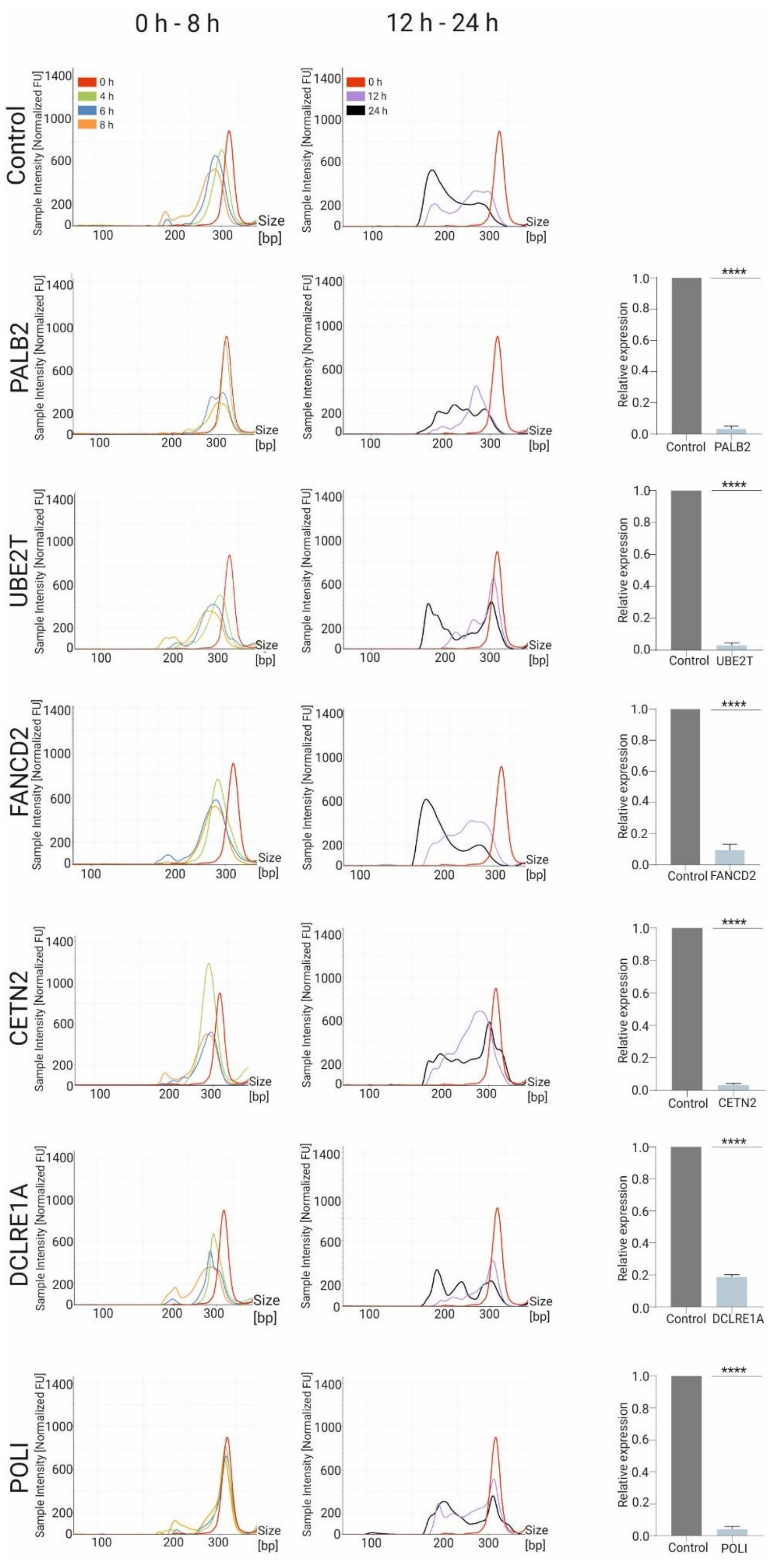


**Figure S7.** Changes in the CAG repeat contraction patterns following the knockdown of selected candidates identified in the CRISPRi screen. dCas9-KRAB-expressing RPE-1 cells were infected with viruses carrying gRNAs targeting the TSS of the control (nontargeting gRNA), PALB2, UBE2T, FANCD2, CETN2, DCLRE1A, and POLI. Infected cells were treated with Cas12a_g14 and collected at early time points (0 h, 4 h, 6 h, and 8 h) and later time points (12 h and 24 h). The knockdown efficiency for the genes selected from the CRISPRi screen is shown on the right. Statistical analysis was performed using an unpaired t-test (n = 3), with $p \leq 0.0001$.

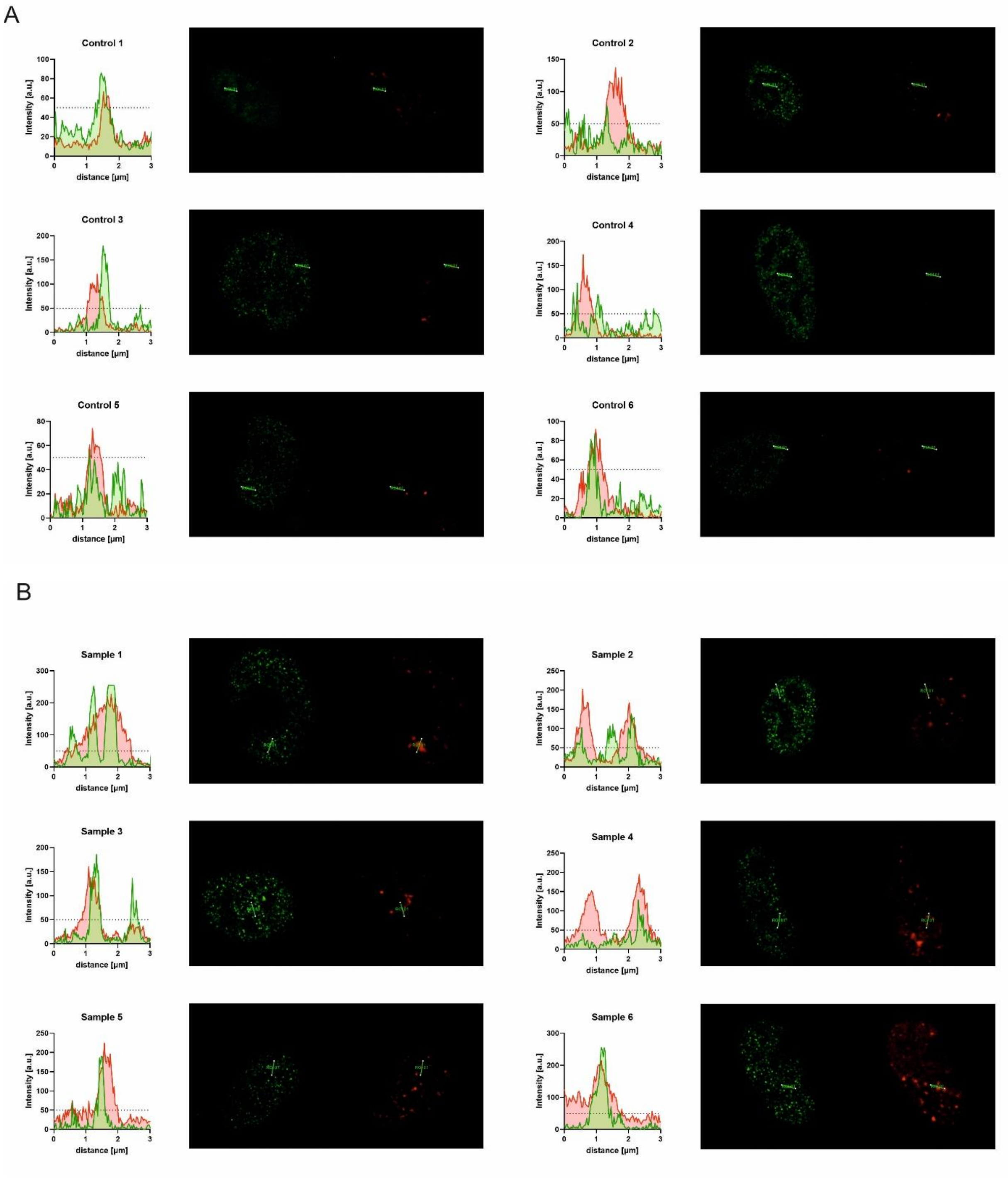


**Figure S8.** Fluorescence colocalization analysis shown as fluorescence line profiles extracted from source micrographs used in the statistical analysis for control **(A)** and cells treated with Cas12a_g14 **(B)**. To visualize the colocalization threshold, a standard cutoff of 50 intensity units was applied. All profiles are 3 µm long and were generated using Leica Application Suite X version 3.4.2.18368 with the "Line profile" module without oversampling.

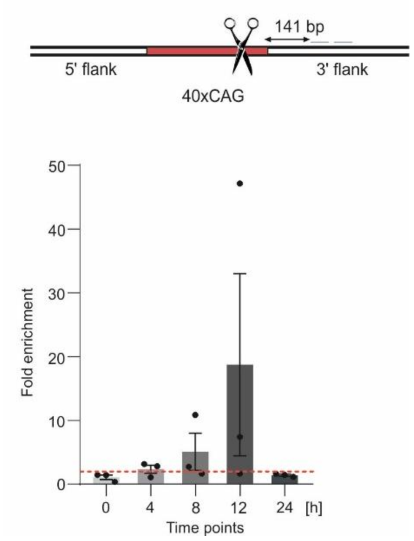


**Figure S9.** ChIP–qPCR analysis of DCLRE1A enrichment at the flanking region of 40 CAG repeats after DSB induction. The enrichment of DCLRE1A was analyzed at 4 h, 8 h, and 12 h. The primers were located on the 3' flank, 141 bp downstream of the repetitive tract. The data shown in this figure represent the means ± SEs (n = 3). A twofold increase in enrichment was considered significant (marked with a red line). No significant differences were detected between protein enrichment at different time points, as indicated by the results of the Kruskal–Wallis test (P value = 0.0556).

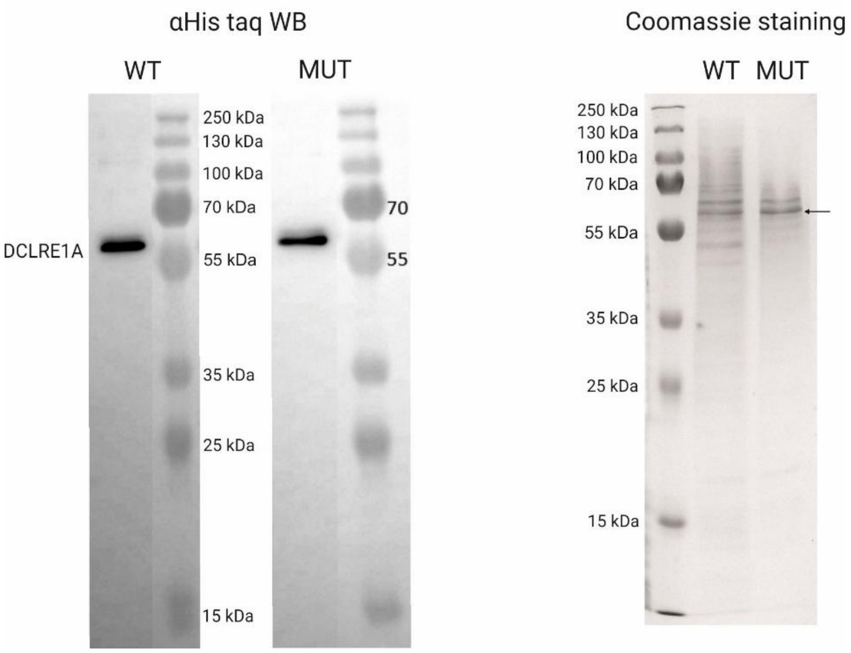


**Figure S10.** Polyacrylamide gel showing the recombinant wtDCLRE1A (WT) and mutant DCLRE1A (MUT) proteins. The PageRuler™ Plus Prestained Protein Ladder was used as a molecular weight marker.

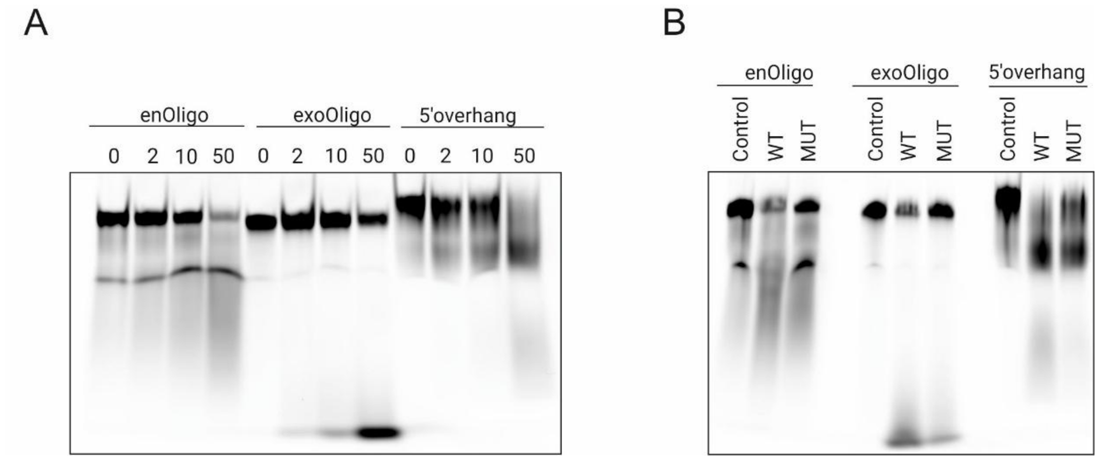


**Figure S11.** Concentration-dependent cleavage activity of recombinant DCLRE1A protein and comparison of cleavage activity of wild-type and mutant variants. **A.** The purified recombinant DCLRE1A protein exhibited endo- and exonucleolytic activities toward structures formed by 5' overhangs in vitro. The DCLRE1A protein (0, 2, 10 and 50 nM) was added to the annealed oligonucleotides (enOligo and exoOligo) and a positive control with a 5' overhang. The digested products were separated on a PAGE gel under denaturing conditions. **B.** Comparison of recombinant wtDCLRE1A and mutDCLRE1A proteins on enOligo and exoOligo, and a positive control with a 5' overhang.

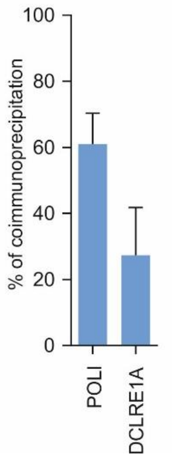


**Figure S12.** Coimmunoprecipitation of Flag-tagged wtDCLRE1A or FLAG-tagged POLI following nitrogen mustard treatment.

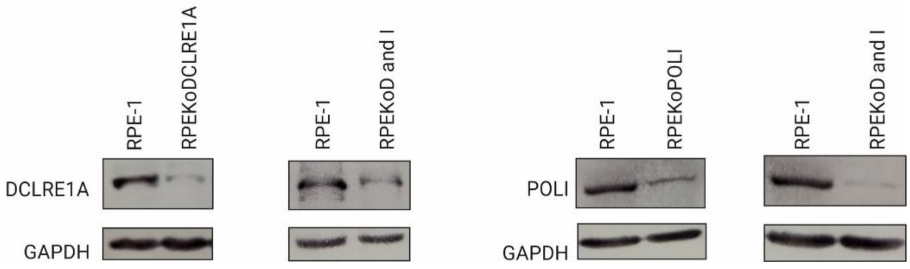


**Figure S13**. Generation of single and double knockout cell lines. Cell lines with a knockout of DCLRE1A (RPEKoDCLRE1A) or POLI (RPEKoPOLI) and a double knockout of DCLRE1A and POLI (RPEKoD and I) were generated using CRISPR-Cas9. The knockout efficiency was confirmed by Western blot analysis.

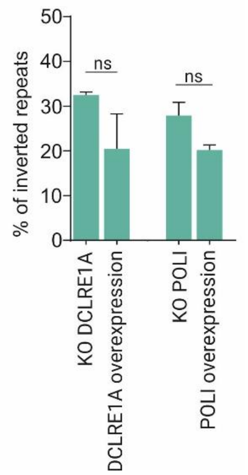


**Figure S14.** The frequency of inverted repeats. Rescue experiments were conducted in RPE1koDCLRE1A and RPEkoPOLI cells following the overexpression of DCLRE1A and POLI, respectively. The data represent two independent biological replicates.

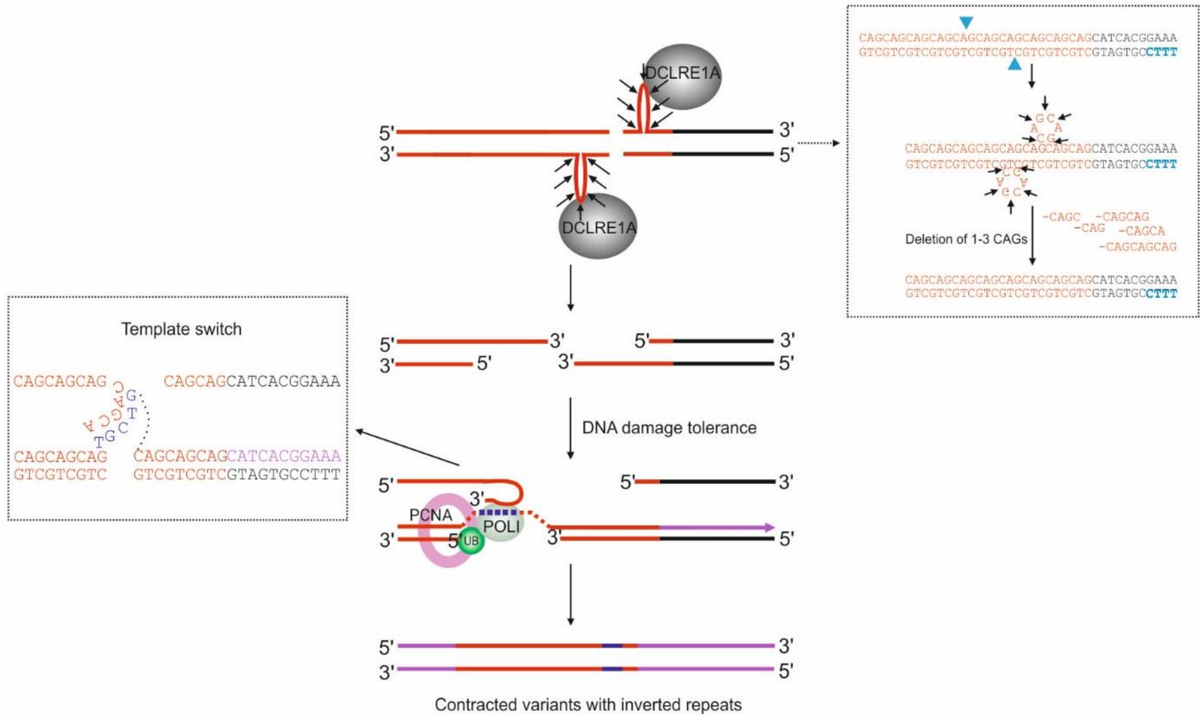


**Figure S15.** Model showing how DCLRE1A and POLI may contribute to the formation of inverted repeats. DCLRE1A is recruited to the secondary structure, where it cuts the heterologous loop, leading to the deletion of one or more CAG repeats. These short deletions expose a 3’ overhang, which promotes strand invasion. POLI subsequently initiates DNA synthesis, which is capable of replicating through regions with secondary structures. This process facilitates a template switching mechanism, resulting in the formation of inverted CAG repeats.

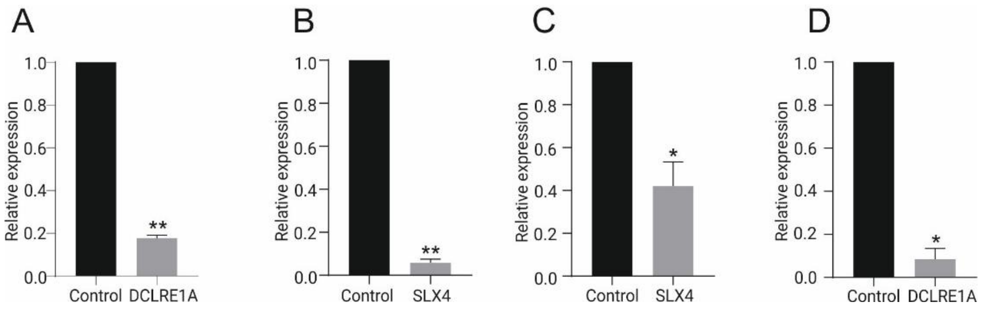


**Figure S16.** CRISPRi-based knockdown of DCLRE1A and SLX4. Graphs **A** and **B** illustrate the knockdown efficiency of DCLRE1A and SLX4 following the transduction of RPE-1 cells with the virus carrying the gRNA complementary to the TSS of DCLRE1A or SLX4. Graphs **C** and **D** depict the knockdown efficiency of both DCLRE1A and SLX4 in cells experiencing simultaneous inhibition of both genes. Statistical analysis was performed on logarithmic values using an unpaired t-test (n=2), where $p \leq 0.0001$.

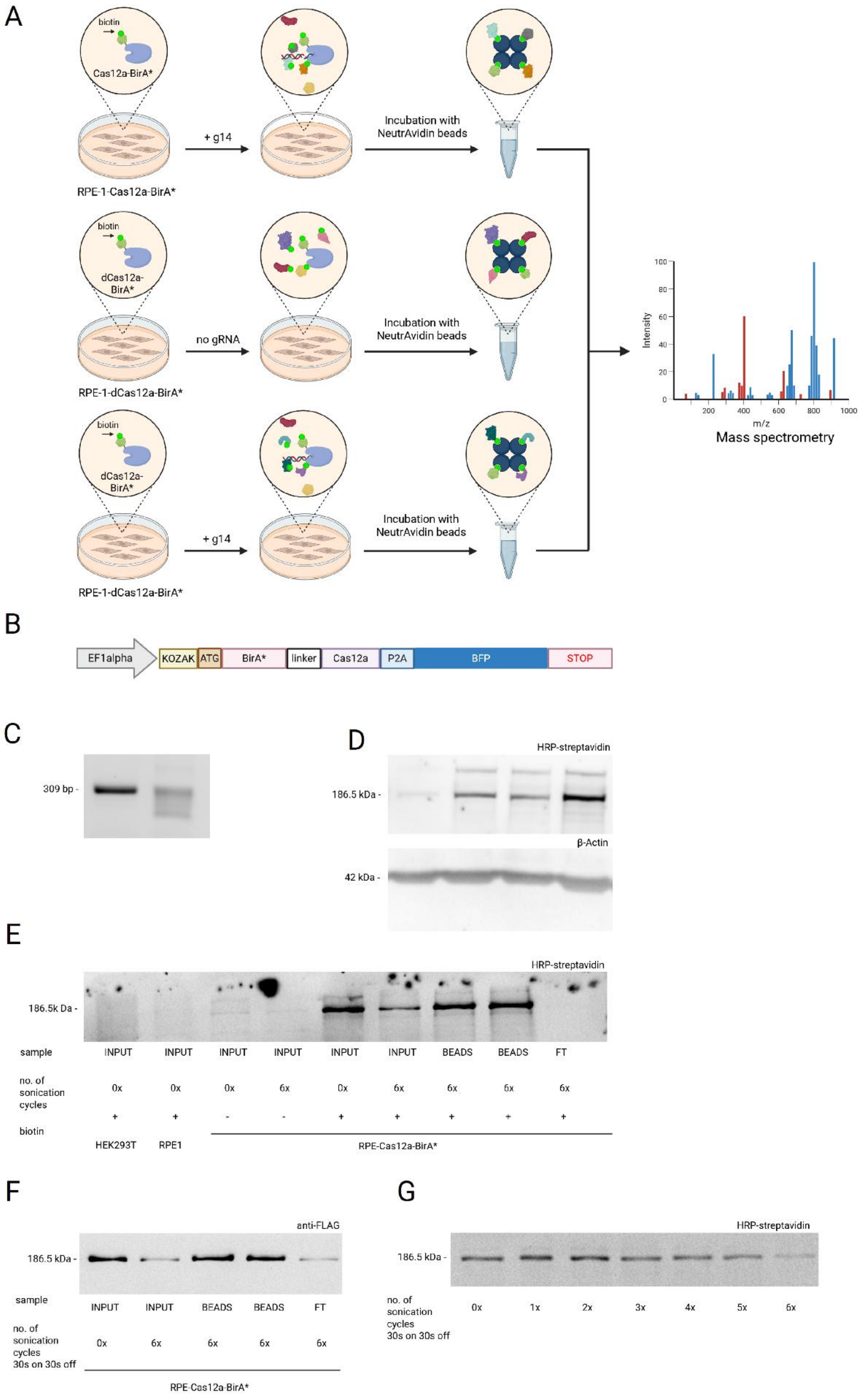


**Figure S17.** BioID analysis. **A.** Schematic representation of the BioID experiment. The RPE1 cell line, constitutively expressing Cas12a-BirA*, was cultured in biotin-supplemented medium for 16 hours and lipofected with g14. Cas12a-BirA* subsequently cleaves the target sequence while the proteins near the target region are labeled with biotin. Cells were collected 12 h after g14 delivery. Labeling with biotin enables pull-down with NeutrAvidin beads and the capture of proteins present at the cleavage site, including DNA repair proteins. The control samples included RPE-dCas12a-BirA* cells lipofected with or without g14. **B.** Schematic representation of the construct used to generate cell lines with stable expression of Cas12a linked to the BirA* domain. RPE-1 cells were transduced with a lentivirus carrying a plasmid with the construct and then sorted based on the BFP signal. C-D. Individual clones were validated for both Cas12a activity and BirA* activity. **C.** Evaluation of Cas12a activity in separate clones. Gradual shortening of the PCR-amplified product after editing by Cas12a following lipofection with g14. **D**. Detection of the biotinylation of Cas12a-BirA* (186.5 kDa), which is the most highly biotinylated protein in cells expressing the construct, after incubation with biotin. The cell lysates were subjected to analysis with HRP-streptavidin. β-Actin was used as a loading control. **E.** Pull down of biotinylated Cas12a-BirA* with NeutrAvidin beads in cell lysates. Detection of biotinylated Cas12a-BirA* with HRP-streptavidin. FT (flow through), sample obtained after the incubation with NeutrAvidin beads; beads, sample eluted from NeutrAvidin beads; input, sample obtained before the incubation with NeutrAvidin beads. **F.** Sonication of Cas12a-BirA* cell lysates. Two cycles of sonication (30 s on and 30 s off) were applied in further experiments.

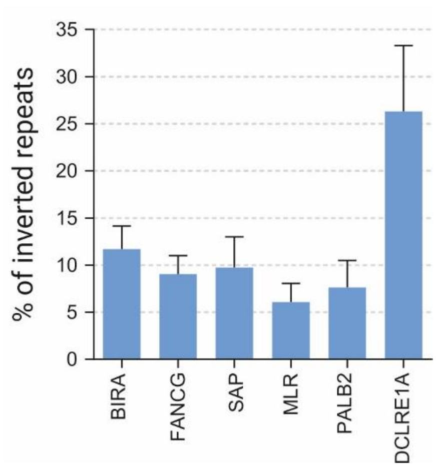


**Figure S18.** Frequency of inverted repeats following Cas12a-DNA repair factor-mediated DSBs within CAG repeats.
